# Autonomous sampling and eDNA metabarcoding for offshore biodiversity monitoring: a pilot study around an FPSO in the Southwestern Atlantic

**DOI:** 10.64898/2026.10.22.761088

**Authors:** Danielle L. A. S. do Amaral, João Gabriel Rodinho Nunes Ferreira, Rosa M. B. Cavalcanti, Marcelo Merten, Juliana Americo, James Birch, Christina Preston, Nadia Allaf, Scott Jensen, Sylvain Joyeux, Italo Marques, Rafael Coelho, Christopher A. Scholin, Mauro F. Rebelo

## Abstract

Characterizing biodiversity around offshore infrastructure is necessary to assess possible influences of industrial activities on marine communities and distinguish persistent ecological change from short-term variability. Combining environmental DNA (eDNA) metabarcoding with autonomous sampling offers a way to expand biological, spatial and temporal coverage while reducing dependence on crewed field operations. We evaluated this approach around a floating production, storage and offloading unit (FPSO) in the Santos Basin, Brazil, to establish an initial biodiversity reference and inform subsequent monitoring. A third-generation Environmental Sample Processor integrated into the W Tupan II uncrewed surface vessel enabled autonomous water sampling at coastal sites, transit locations and the FPSO area, yielding 64 biological samples. We used mitochondrial 12S and 16S metabarcoding to assess taxonomic diversity, sampling completeness and spatial and short-term temporal variation. Around the FPSO, 47 samples from 21 points detected 34 species with 12S and 28 with 16S, representing 74.3% and 65.4% of the respective estimated species richness. Local analyses showed weak radial differentiation across 300–1,500 m around the FPSO, with no near–far taxonomic indicators remaining significant after multiple-testing corrections. Samples collected at the same point approximately 1–2 h apart were almost as dissimilar as samples from different points, whereas technical replicates of post-extraction processing closely matched their source DNA profiles. Taxon turnover accounted for approximately 73% and 80% of temporal Jaccard dissimilarity for 12S and 16S, respectively. Regional analyses recovered a coastal–offshore compositional contrast, with geography and sampling date jointly represented. This pilot supports autonomous eDNA sampling as an operational approach to offshore biodiversity monitoring. The findings favour repeated sampling across campaigns to distinguish transient detections from recurrent patterns and persistent compositional changes.

## Introduction

Biodiversity monitoring around offshore infrastructure requires observations that can distinguish persistent ecological change from the variability of marine assemblages. Obtaining these observations is difficult when surveys depend on crewed vessel availability, specialist personnel and discrete field campaigns as the biological community and associated water masses continually evolve between visits. Expanding spatial coverage alone does not resolve this mismatch: repeated observations are also needed to establish which components of an assemblage recur and which detections reflect shorter-lived conditions. The logistical constraints of marine sampling and the temporal variability documented in coastal surveys therefore motivate monitoring approaches that combine broad biological coverage with repeatable sampling [1– 3].

Environmental DNA (eDNA) metabarcoding combines DNA-based taxonomic identification with high-throughput sequencing of genetic markers recovered directly from environmental samples [4, 5]. In seawater, this DNA pool includes material within captured microorganisms and small metazoans, as well as extra-organismal DNA released by larger organisms [5]. Previous work has shown that multimarker surveys resolve spatial structure in benthic, planktonic and fish assemblages around offshore infrastructure, while repeated sampling reveals changes in community composition and taxon recurrence across temporal scales [2, 3, 6, 7].

The spatial and temporal resolution of marine eDNA depends on the interaction between biological inputs, hydrodynamic transport, settling and degradation. Advection can displace DNA from its source, while settling and decay constrain its distribution and persistence; the resulting spatial footprint depends on environmental conditions and the properties of the sampled material [8]. Marker selection and sequencing depth further affect which taxa are recovered [9]. Temporal observations have revealed diel changes, seasonal shifts and interannual recurrence in marine eDNA assemblages, demonstrating that the timing and frequency of sampling influence the community represented [2, 3, 10]. These findings make temporal replication relevant to offshore monitoring, where single-timepoint observations can describe transient fish distributions without establishing persistent associations with infrastructure [7].

Autonomous underwater vehicles and uncrewed surface vessels offer complementary ways to extend eDNA sampling beyond the spatial and temporal coverage achievable through intermittent crewed campaigns. Integrating these vehicles with the Environmental Sample Processor (ESP), which automates water filtration and sample preservation, has enabled repeated sampling of water masses, multi-marker biodiversity surveys and sample acquisition along long-distance open-ocean transects \cite{ref1,ref11,ref12}. Coupling autonomous sampling with high-throughput sequencing provides a scalable framework for collecting and analysing large numbers of samples at the frequency and spatial coverage required for biodiversity monitoring across extensive offshore areas \cite{ref11,ref12}. The combination of autonomous mobility and on-board sample processing can reduce dependence on dedicated crewed vessel time and manual field operations, addressing logistical and cost constraints on sustained sampling \cite{ref1,ref12}. Remote operation also provides a means of reducing the exposure of sampling personnel to offshore hazards, while retaining the need to manage equipment and operational risks \cite{ref13}.

Offshore monitoring is needed to characterize biodiversity and assess possible influences of industrial activities and structures on marine communities. Molecular surveys can support this assessment by identifying compositional differences and relating them to environmental conditions and reference areas \cite{ref6,ref7,ref14}. Analyses of sampling effort and community composition can help evaluate what an eDNA survey has represented, as illustrated by a large-reservoir study that combined multiple markers, rarefaction and multivariate comparisons \cite{ref15}.

In this pilot study, we evaluated an autonomous sampling system coupled with eDNA metabarcoding as an operational approach to biodiversity monitoring around a floating production, storage and offloading unit (FPSO) in the Santos Basin off southeastern Brazil. A third-generation ESP installed on the W Tupan II uncrewed surface vessel [16] supported a campaign spanning coastal locations, transit waters and the FPSO sampling area. Using mitochondrial 12S and 16S rRNA gene markers, we characterized biodiversity and spatial and shortterm temporal variation in community composition. The objective was to establish an initial site characterization and identify requirements for subsequent monitoring of offshore biodiversity using an uncrewed platform.

## Materials and methods

### Study area, campaign and sampling design

The study was conducted around a floating production, storage and offloading unit (FPSO) in the Santos Basin off southeastern Brazil, and along a coastal-to-offshore route connecting the offshore sampling area to Rio de Janeiro (Fig. 1). The FPSO began operating in October 2025, approximately six months before the sampling campaign. Coastal sampling included Ilhas Cagarras and Urca, near the entrance to Guanabara Bay, Rio de Janeiro. The campaign comprised 64 biological samples collected from 1–17 April 2026: 47 around the FPSO, nine during the return transit, six from Ilhas Cagarras, and two from Urca (Table 1).

**Table 1.** Sampling groups, independent biological samples, geographic points, and sampling dates.

| Group | Locality | Biological samples (n) | Geographic points (n) | Sampling date(s), UTC |
| --- | --- | --- | --- | --- |
| Coastal | Ilhas Cagaras | 6 | 1 | 1 April 2026 |
| Coastal | Urca | 2 | 1 | 6 and 8 April 2026 |
| Transit | Return transit | 9 | 9 | 16–17 April 2026 |
| FPSO | FPSO area | 47 | 21 | 13–15 April 2026 |
| Total | — | 64 | 32 | 1–17 April 2026 |

**Fig. 1.**
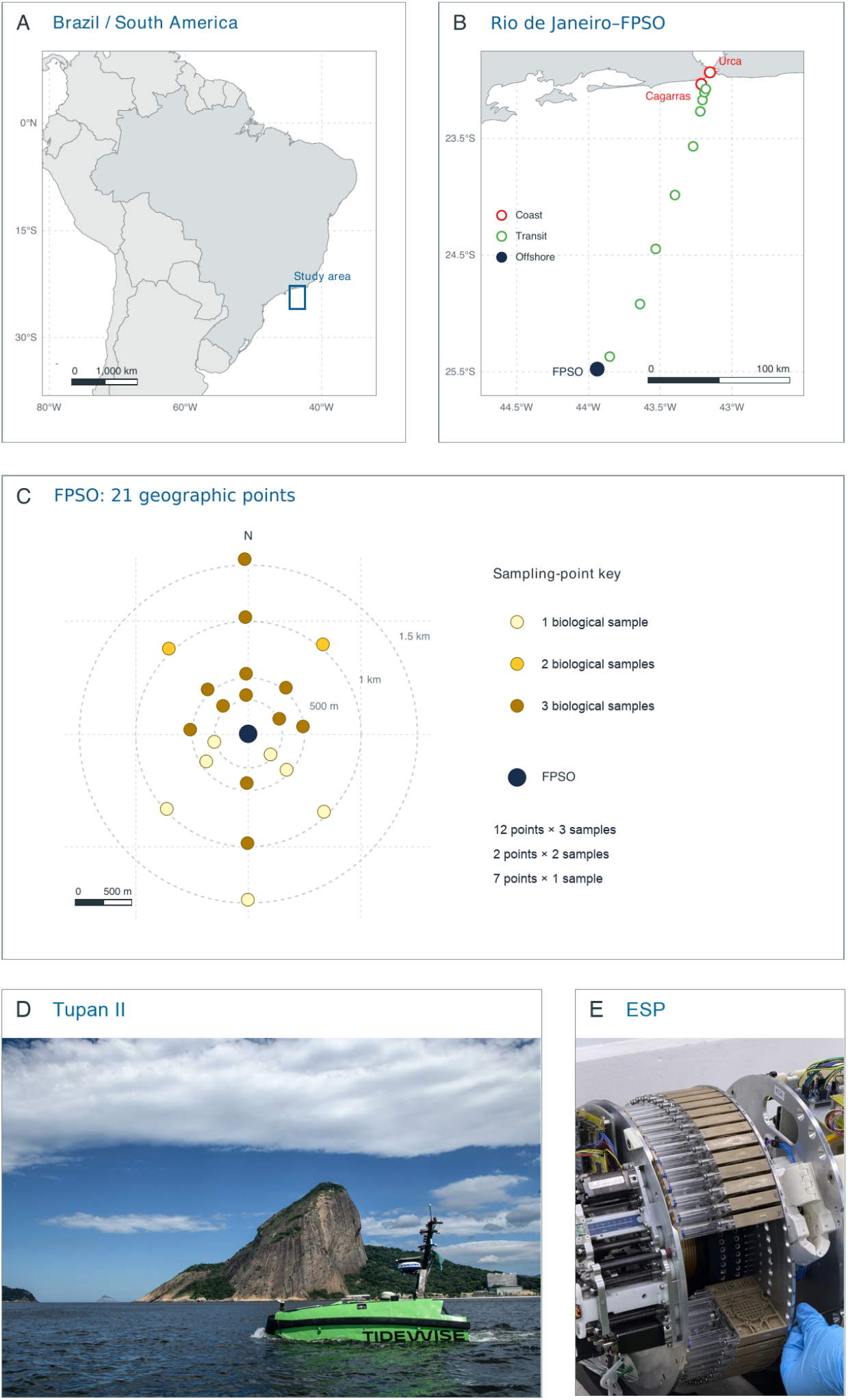
Sampling design and autonomous sampling platform of the FPSO eDNA campaign. The campaign comprised 64 biological water samples at 32 geographic points: 47 samples at 21 points around the FPSO, nine samples along the return transit, six at Ilhas Cagarras, and two at Urca, Rio de Janeiro, Brazil. **A**. South American locator map; the blue rectangle marks the study area off southeastern Brazil. **B**. Coastal-to-offshore sampling corridor. Open red circles identify the coastal locations, open green circles identify transit locations, and the filled dark symbol marks the offshore FPSO area, whose individual points are resolved in C. **C**. Local sampling points relative to the FPSO (larger filled dark circle). Smaller filled yellow circles of equal size identify geographic points; progressively darker yellow indicates one, two or three biological samples. Twelve points were sampled three times, two twice, and seven once. Dashed rings indicate distances of 300, 500, 1,000 and 1,500 m from the FPSO. North is upward. A–B use longitude–latitude maps (plate carrée), whereas C uses local east–north distances. Scale bars indicate 1,000 km, 100 km and 500 m in A, B and C, respectively. Coastlines are from Natural Earth at 1:50 million resolution in A and 1:10 million in B. **D**. Side view of Tupan II traveling parallel to the coast. Sugarloaf Mountain dominates the background, with buildings visible near the shoreline. Image courtesy of Luciano Candisani/Bio Bureau. **E**. Environmental Sample Processor (ESP), developed by the Monterey Bay Aquarium Research Institute (MBARI), showing its sampling-cartridge assembly. The ESP automates water filtration and sample preservation for environmental DNA (eDNA)-based surveys of marine biodiversity. Image courtesy of Nadia Allaf/MBARI.

Offshore sampling was performed using a third-generation Environmental Sample Processor (3G-ESP, Monterey Bay Aquarium Research Institute, USA) integrated into the W Tupan II unmanned surface vessel (TideWise, Brazil). The vessel and the ESP were operated remotely from TideWise’s Remote Operations Center in Rio de Janeiro. The local sampling grid incorporated stations from Equinor’s environmental monitoring programme and additional locations at planned radial distances of 300, 500, 1,000 and 1,500 m from the FPSO. Sampling locations and their execution order were adjusted to accommodate security access restrictions.

The 64 biological samples were grouped by sampling location: FPSO (47 samples), coastal (eight samples) and transit (nine samples) (Table 1). Each represented a separate water sampling and filtration event.

FPSO sampling took place on 13–15 April 2026 at 21 geographic points. Twelve points were sampled three times, two points twice, and seven points once (Fig. 1). Repeated sampling at the same point was conducted at different times to assess short-term variability.

Coastal sampling comprised six biological samples collected at Ilhas Cagarras on 1 April 2026 and two at Urca on 6 and 8 April 2026 during system testing. Nine biological samples were collected during the return transit on 16–17 April 2026. Regional analyses included all three sampling groups; local analyses included only the FPSO samples.

Libraries were prepared manually for all biological samples. To compare library preparation methods, we selected 11 samples, one from each of the 11 points in FPSO’s traditional monitoring grid. For each selected sample, an additional technical library preparation used the same DNA aliquot. Only clean-up was automated on an Opentrons OT-2 using the Clean-up 2 protocol; reaction assembly and indexing were manual. We compared manually cleaned and robot-cleaned preparations as 11 matched pairs. These additional libraries were technical replicates, not additional biological samples, and were used only for the method comparison, not for richness estimates or ecological models.

### Filtration and DNA extraction

Water was pumped and filtered autonomously in the field by the third-generation Environmental Sample Processor (3GESP; MBARI, USA) installed on the W Tupan II uncrewed surface vessel. The mechanical design and function of the 3G ESP have been previously described [17]. Sampling cartridges contained polyvinylidene fluoride (PVDF) membranes with a pore size of 0.22 µm and a diameter of 25 mm (Durapore, Millipore; GVWP02500). The ESP was programmed to filter up to 1 L per sample, stopping earlier if the filter clogged, with a maximum filtration time of 1 h. The filtered volume was recorded automatically for each sampling event. Filtered volumes for individual samples are provided in Supplementary Data 1. After filtration, retained material was preserved with RNAlater (Sigma) inside the cartridge. At the end of sampling, the RNAlater was removed by the ESP. Filters were frozen at −80 °C without washing and stored until DNA extraction. Three field filtration blanks (PR347D, PR406D and PR534D) accompanied the environmental samples. PR534 was prepared after PR406 failed to reach the intended 1 L filtration volume, to verify system operation after the problem was corrected. The laboratory input lists for each marker comprised 64 environmental DNAs and these three blanks.

DNA was extracted using the DNeasy Blood & Tissue Kit (Qiagen), with modifications for filter membranes. Forceps and scissors were flame-sterilized between samples. One half was used for extraction and the other remained frozen at −80 °C as a reserve. The membrane was not washed after thawing or before lysis. The extraction half was incubated overnight at 56 °C and 1,000 rpm in buffer ATL and Proteinase K. After vortex mixing and centrifugation to remove solid particles, buffer AL and ethanol were added to the lysate. The mixture was loaded onto DNeasy Mini Spin columns and washed with buffers AW1 and AW2 according to the manufacturer’s instructions. DNA was eluted in buffer AE preheated to 70 °C. The eluate was reapplied to the same column, incubated for 10 min at room temperature and recovered by a second elution. The DNA concentration was determined using the Qubit dsDNA HS Assay Kit (Thermo Fisher Scientific). DNA extracts were stored at −80 °C.

### Library preparation and sequencing

The mitochondrial 12S and 16S rRNA markers were amplified using MiFish-U [18] and MarVer3 [19] primers, respectively. For 12S, amplification used two PCR steps before indexing. The first used primers without overhangs, Platinum SuperFi II PCR Master Mix (Thermo Fisher Scientific) and 25 µg bovine serum albumin (BSA; Thermo Scientific, B14; stock 20 mg/mL; final concentration 1 mg/mL). Reaction compositions, primer sequence and concentrations and thermocycling conditions are provided in Supplementary Data 2.

The second 12S PCR added Illumina overhangs using Platinum II Hot-Start PCR Master Mix (Thermo Fisher Scientific), Platinum GC Enhancer, MiFish-U forward and reverse primers carrying Illumina overhangs, and the first PCR product.

For 16S, MarVer3 primers carrying Illumina overhangs were used in a single amplification step before indexing. Each 25 µL reaction used Platinum SuperFi II PCR Master Mix and 25 µg BSA (final concentration 1 mg/mL), as detailed in Supplementary Data 2.

PCR products were purified with Illumina Purification Beads. Indexing reactions (50 µL) used DreamTaq PCR Master Mix (Thermo Fisher Scientific), 50 µg BSA (final concentration 1 mg/mL) and precombined Illumina DNA/RNA UD Indexes, Tagmentation (i5/i7). Index sets C (20091658) and D (20091660) were used for 12S and set A (20091654) for 16S. Indexing reactions were assembled manually. Indexed products underwent a second purification with Illumina Purification Beads. Reagent quantities and bead ratios are provided in Supplementary Data 2.

For 11 selected FPSO samples, additional technical preparations underwent clean-up on an Opentrons OT-2 using the Clean-up 2 protocol and were compared with manually cleaned preparations from the same source DNA. Only cleanup was automated; reaction assembly and indexing were manual. These technical preparations were assessed separately from biological sampling variation.

Amplification and indexing were assessed by electrophoresis on 3% agarose gels, and representative libraries were examined using an Agilent TapeStation. Final libraries were quantified with the Qubit High Sensitivity system (Thermo Fisher Scientific), normalized as specified in Supplementary Data 2 and sequenced on an Illumina NovaSeq X Plus with the Illumina NovaSeq X Series 1.5B Reagent Kit (300 cycles), using paired-end 150 bp reads (2 × 150 bp).

### Bioinformatic processing and taxonomic assignment

#### Read processing and OTU inference

Paired-end reads from the 12S MiFish and 16S MarVer3 assays were processed independently with the Todabio pipeline, using Nextflow 26.04.4 on AWS Batch and a containerized QIIME 2 2023.9 environment [20]. Read-pair and sample-manifest correspondence was verified, and FastQC and MultiQC reports were inspected for sequencing depth, quality, adapter content and read lengths.

Demultiplexed reads were imported in Casava single-lane format. Cutadapt 4.5 [21] removed the MiFish and MarVer3 primers described in Section 2.3, using a maximum error rate of 0.10, a minimum overlap of 3 bp and a minimum retained length of 1 bp, without additional quality trimming. Reads were then quality-filtered, denoised, dereplicated, merged and screened for chimeras with q2-dada2 [22]. No fixed-length truncation was applied in either direction. Other settings followed the q2-dada2 defaults: two maximum expected errors per read, quality truncation at Phred 2, a minimum paired-read overlap of 12 bp, independent sample processing and consensus chimera removal. Nonchimeric representative sequences were screened against the pipeline’s UniVec reference to remove vector and other technical contaminants.

DADA2-derived amplicon sequence variants were clustered de novo at 99% nucleotide identity within each library. Feature tables and representative sequences were then merged by marker and reclustered at 99% identity to define OTUs consistently across all samples. Centroid sequences represented OTUs, and matrix entries recorded read counts per library. Per-library processing counts were retained for Supplementary Data 3a–b.

#### Taxonomic assignment

OTU representative sequences were queried with BLAST+ against the locus-specific MIDORI2 GB269 references [23]: mitochondrial small-subunit rRNA for 12S and large-subunit rRNA for 16S. Searches required at least 70% nucleotide identity and 50% query coverage and retained up to 50 hits per OTU. Sequences without a MIDORI match were searched against the NCBI nucleotide database snapshot of 8 July 2026, using the same thresholds and up to 25 target sequences per query.

MIDORI lineages and NCBI taxonomy records were used to assign OTUs to the most informative available canonical rank, from domain to species. OTUs without an informative canonical-rank assignment were excluded from taxon-level tables; counts from OTUs sharing an assignment were summed within each library (marker).

#### Taxon-table decontamination

Taxonomic decontamination used the reverse-osmosis water controls PR347D, PR406D and PR534D, while preserving the original tables. Candidate screening considered occurrence in at least two controls, prevalence in no more than 5% of environmental libraries, and an ecological blocklist of human, livestock, domestic-animal and terrestrial taxa. Ecologically plausible marine taxa were protected from automatic removal. Candidates were manually reviewed using prevalence and relative abundance in controls and environmental libraries.

Seven taxa per marker and the three control columns were removed, yielding 105 12S and 132 16S taxonomic assignments across 75 library columns before subsequent taxonomic curation. A separate author-reviewed step reduced five species-level assignments to genus level. In 12S, reads assigned to *Brevoortia tyrannus* and *Cubiceps paradoxus* were merged into the existing *Brevoortia* and *Cubiceps* taxa, respectively. In 16S, *Exocoetus monocirrhus* was merged into *Exocoetus*, and *Paracentrotus lividus* and *Luidia magnifica* were relabelled as *Paracentrotus* and *Luidia*. This conservative reduction in resolution did not assign an alternative species or change per-library read totals. The final curated matrices contained 103 12S and 131 16S assignments (Supplementary Data 4a–b). Matching curated labels between markers produced a presence-only inventory of 199 assignments, including 35 shared between markers. Raw read counts were not summed across loci for this inventory. The combined normalized representation used for ecological analyses is described separately in Section 2.5.

#### Target-locus curation and final OTU matrices

OTUs matching the expected MIDORI locus and measuring at least 100 bp were retained as on-target. OTUs without a MIDORI match were evaluated within operational length windows of 150–200 bp for 12S and 200–300 bp for 16S, provided that no residual primer signature occurred near either terminus. They were retained when their best NT matches indicated mitochondrial or ribosomal RNA and the lowest common ancestor of hits within two percentage points of the best identity remained within Eukaryota. Sequences with insufficient reference support, incompatible lengths, residual primer signatures or non-target matches were excluded.

Target-locus filtering retained 225 of 262 12S OTUs and 253 of 688 16S OTUs. OTUs whose maximum relative abundance in controls exceeded that in environmental libraries were proposed as contaminants and manually reviewed. Removing four OTUs per marker and the control columns left 221 12S and 249 16S OTUs. A subsequent sequence-specific review excluded nine additional 12S and eight additional 16S OTUs as probable contaminants, yielding the database-aligned matrices used for downstream analyses: 212 and 241 OTUs, respectively, across 75 library columns (Supplementary Data 5a–b). These columns represent 64 biological samples and 11 technical preparations, whose analytical inclusion is defined in Section 2.5.

### Analytical datasets, sample inclusion and transformations

Analyses used the marker-specific taxon and OTU count matrices described in Section 2.4, linked to sampling metadata through the sampling, filtration and DNA records. We used in-house Python scripts (`extrai-matrizes.py` and `09b-prepara.py`) to assemble sample-by-unit matrices, harmonize identifiers and derive analytical metadata. Taxa retained their assigned resolution; an unresolved genus or family was not treated as the sum of its identified descendants. OTUs remained separate molecular units even when they shared a taxonomic assignment.

The regional dataset comprised 64 biological samples, including 47 samples at 21 FPSO points. One manually prepared library profile per sample was retained for ecological analyses. The 11 additional Opentrons preparations were used only for technical comparisons. Descriptive local composition and incidence-based richness analyses retained all 47 biological samples. Multivariate analyses retained libraries with at least 1,000 reads in the corresponding taxon or OTU matrix, removed units with zero counts in the retained subset, and required the metadata relevant to each model. The resulting regional sample sizes were 63 for 12S taxa and OTUs and 62 for 16S taxa and OTUs. Local taxonomic comparisons retained 46 biological profiles for 12S and 45 for 16S; full local models further required complete predictor data and used 43 samples per marker.

Within each sample, counts were divided by the total for the corresponding representation. Hellinger-transformed profiles were obtained by taking the square root of these proportions, and compositional dissimilarity was calculated as Euclidean distance between transformed profiles. The combined regional taxonomic representation matched taxon identities across markers and summed their separately normalized profiles before Hellinger transformation, giving equal weight to each nonempty marker profile. A marker with no assigned reads contributed zero. Its inclusion filter used the sum of taxonomic read counts across markers, rather than requiring 1,000 reads separately in each marker; all 64 biological samples passed this combined filter. Relative read abundance was used as a molecular compositional measure, not an estimate of organism abundance.

Statistical analyses were implemented in R and in-house Python scripts. The recorded project toolchain comprised R 4.5.2, vegan 2.6.10 and Python 3.14.4. In-house Python routines used NumPy for matrix operations, estimators and permutation algorithms, with Matplotlib for graphical outputs; the editable Fig. 2 source records Matplotlib 3.9.4. Selective post-curation recalculations used R 4.6.1 with vegan 2.6.10 and Python 3.12 with NumPy 2.0.2 and Matplotlib 3.9.4; the original estimators, cohorts and resampling settings were retained. Count-based OTU analyses and the numerically unchanged local 12S complete-case analyses were retained. Analysis-specific package versions are identified where available. Analysis-specific resampling settings are given below.

**Fig. 2.**
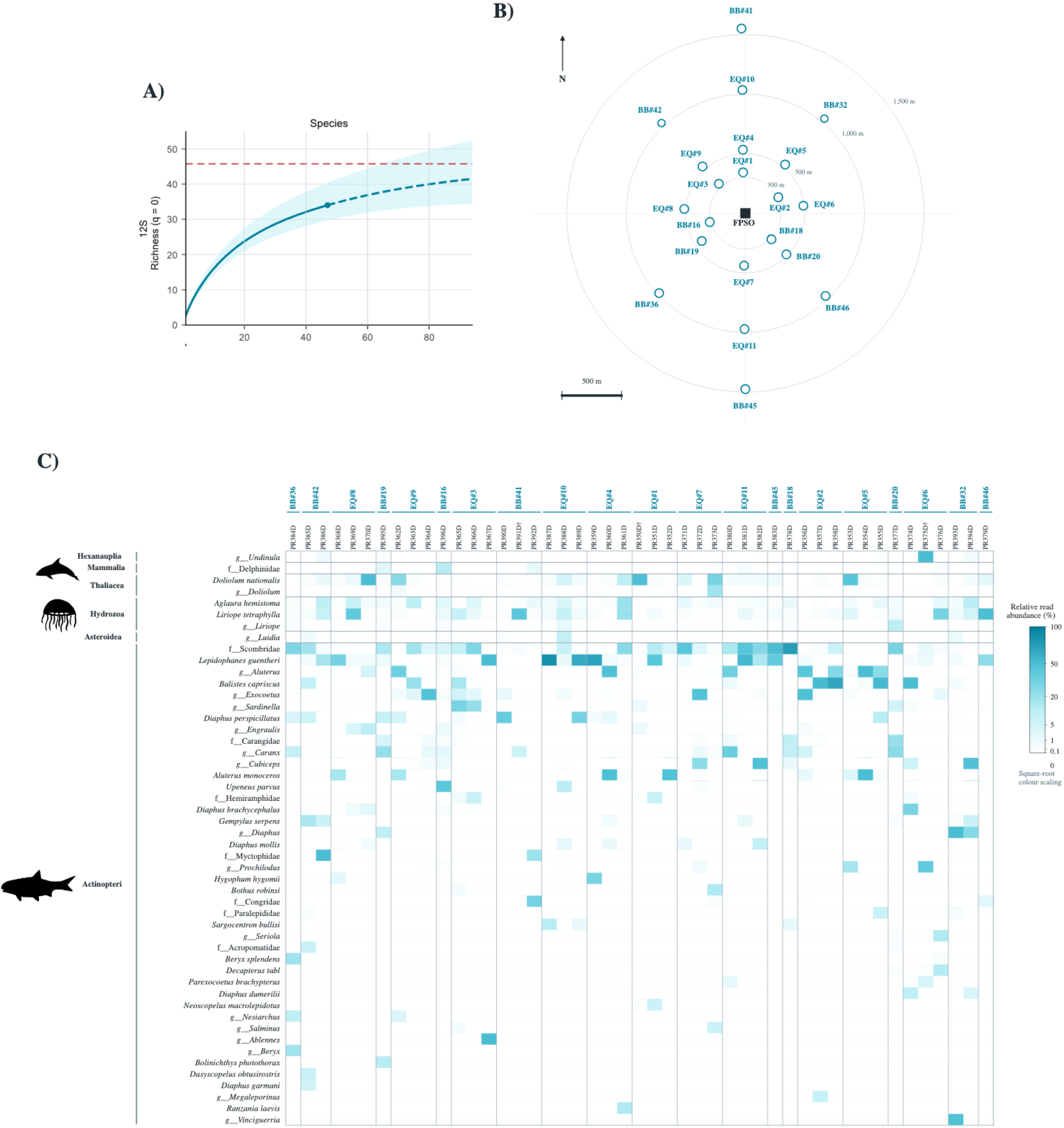
Sampling effort and spatially referenced taxonomic profiles around the FPSO. **A**. Incidence-based richness for 12S species (Hill number q = 0, which counts species without abundance weighting). The horizontal axis represents the number of biological samples. The solid cyan line shows rarefaction up to the observed effort of 47 samples and the dashed segment extrapolates to 94. Cyan shading indicates uncertainty around the curve, and the horizontal dashed red line marks the Chao2 richness estimate. **B**. The 21 sampling points (open cyan circles), FPSO position (dark square), north arrow, 500-m scale bar and distance rings at 300, 500, 1,000 and 1,500 m. Point IDs link the map to the column groups in C. **C**, relative read abundance of the 50 family-, genus- or species-level taxa with the highest mean normalized abundance across the 47 samples. Columns are individual samples, grouped by geographic point from west to east and ordered chronologically within each point. Rows are grouped by taxonomic class, with Actinopteri last, and ordered within each class by decreasing prevalence (number of samples with a nonzero detection). The prefixes f and g denote family- and genus-level assignments; unprefixed names denote species. Daggers mark low-read marker libraries: PR350D and PR375D for 16S, and PR391D for 12S. These samples remain in this descriptive display.

### Local taxonomic diversity and sampling completeness

#### Taxonomic composition and spatially referenced profiles

We described local composition using all 47 standard-protocol profiles, retaining low-read libraries for this descriptive endpoint. The in-house Python script `figure_map_ heatmap_draft_v3.py` combined the two within-marker relative-read profiles and normalized the resulting profile over all assigned taxa before selecting taxa for display. Each marker therefore contributed equally, irrespective of its sequencing depth. Among nonzero species-, genusand familylevel assignments, we selected the 50 taxa with the greatest mean normalized relative read abundance across samples. This subset was not renormalized. Prevalence was the number of samples with a nonzero detection, rather than mean read abundance.

Columns in Fig. 2C were grouped by geographic point from west to east and ordered chronologically within points. Rows were grouped by class, with Actinopteri last, and ordered within classes by decreasing prevalence. Prefixes f and g denote unresolved familyand genus-level taxa, not totals over identified descendants. PR350D and PR375D for 16S and PR391D for 12S were flagged as low-read libraries but retained in the display.

Complete composition by class was calculated separately with the in-house script `local_taxonomic_class_ composition.py` (Supplementary Table S1). We counted distinct detected assignment IDs at their original ranks, assigned them to their recorded class, and retained class-unresolved assignments explicitly. Combined taxon counts were the union of IDs across markers. Raw assigned reads were summed across samples and, for the pooled summary, across markers; percentages used all assigned reads in the corresponding marker or pooled dataset. These read-weighted summaries differ from the equal-marker weighting of the heatmap and are not species-richness estimates.

#### Incidence-based richness, rarefaction and extrapolation

Species richness and OTU richness were analysed separately for each marker using incidence across the 47 biological samples. Species came from species-level taxonomic matrices; multiple OTUs assigned to the same species remained distinct molecular units. Presence required at least one read. Sampling effort counted biological samples, including repeated samples at the same point.

The in-house Python script `figure5_sampling_ sufficiency_draft.py` implemented incidence-based richness estimation (Hill number q = 0), sample-based rarefaction through 47 samples and extrapolation to 94. Its estimator followed the incidence formulation used by iNEXT 3.0.2, which also served as the reference for validation of the project’s in-house R implementation. If T is the number of samples and Q1 and Q2 are the numbers of units detected in exactly one and two samples, estimated unseen richness was [(T − 1)/T] × Q1^2^/(2Q2) when Q2 > 0, and [(T − 1)/T] × Q1(Q1 − 1)/2 otherwise. The Chao2 estimate was observed richness plus this unseen component. Interpolation used the hypergeometric probability of detecting each observed unit; extrapolation approached the Chao2 estimate using the observed increment between T − 1 and T samples. Observed richness divided by Chao2 quantified the proportion of estimated richness detected.

Uncertainty was estimated by 200 parametric bootstrap replicates. Detection probabilities were estimated for observed and unseen units following the iNEXT 3.0.2 incidence-bootstrap procedure; replicate incidence frequencies were drawn from binomial distributions with T trials. At each effort, the interval was the point estimate ± 1.96 bootstrap standard errors, with its lower limit bounded at zero. Fixed seeds derived from 42 were used: 42–45 for the complete species/OTU series, 142–145 for their recurrent subsets, 242– 245 for fish-only series and 342–345 for fish-only recurrent subsets.

Singletons and doubletons were defined by occurrence in one or two samples, not by read counts. A sensitivity analysis excluded units detected in only one of the full 47 samples, fixing subset membership before estimation and retaining all sample rows. This filter sets Q1 to zero and therefore removes the Chao2 unseen-richness component. Fish-only analyses retained lineages in Actinopteri or Chondrichthyes and excluded OTUs without an established fish assignment. Bootstrap intervals treated assignments and subset membership as fixed and did not account for dependence among repeated field samples.

### Local spatial structure and near–far indicator analysis

We fitted marker-specific distance-based redundancy analyses (dbRDA) to Hellinger-transformed taxonomic profiles using Euclidean distances. The full local models used 43 biological samples per marker with at least 1,000 assigned reads and complete sampling-design, current and wind data. For 12S, PR350D, PR351D, PR390D and PR391D were excluded from the 47-sample local cohort; for 16S, exclusions were PR350D, PR351D, PR375D and PR390D. These exclusions reflected the marker-specific read filter and missing wind or current records.

Planned radius entered as a continuous predictor in metres; relative position was a four-level factor (N, S, E, W) derived from bearing relative to the FPSO. sampling hour was recorded in decimal local time (UTC − 3), and the period was morning (06:00–11:59), afternoon (12:00–17:59) or night (18:00–05:59). Current and wind directions were represented by sine and cosine of their bearings, with speeds in knots.

The full additive model was composition ∼ radius + position + hour + period + current_sin + current_cos + current_speed + wind_sin + wind_cos + wind_speed.

In-house R scripts `09e-escores.R` and `10c-misto-aninhado.R` fitted the full models and compared them with design-only models containing radius, position, hour and period on the same complete-case samples. Model and nested-model tests used 999 unrestricted permutations and seed 42. We reported pseudo-F, permutation p, adjusted R^2^ and axis percentages relative to total inertia. Conditional block importance was calculated with `10d-importancia-biplot.R` as the full-model adjusted R^2^ minus that obtained after removing each block while retaining all other blocks and the same observations. Current and wind each formed a block containing their two directional terms and speed. These changes are conditional losses, not additive variance fractions; the complement of adjusted R^2^ is variation not captured by the model.

The in-house Python script `local_community_ ordination.py` computed unconstrained PCoA for the identical 43 profiles per marker through singular-value decomposition of the centred Hellinger matrix. The radius-only and radius-plus-position displays reused the same coordinates (Fig. 3E–H). Axis percentages used total inertia.

**Fig. 3.**
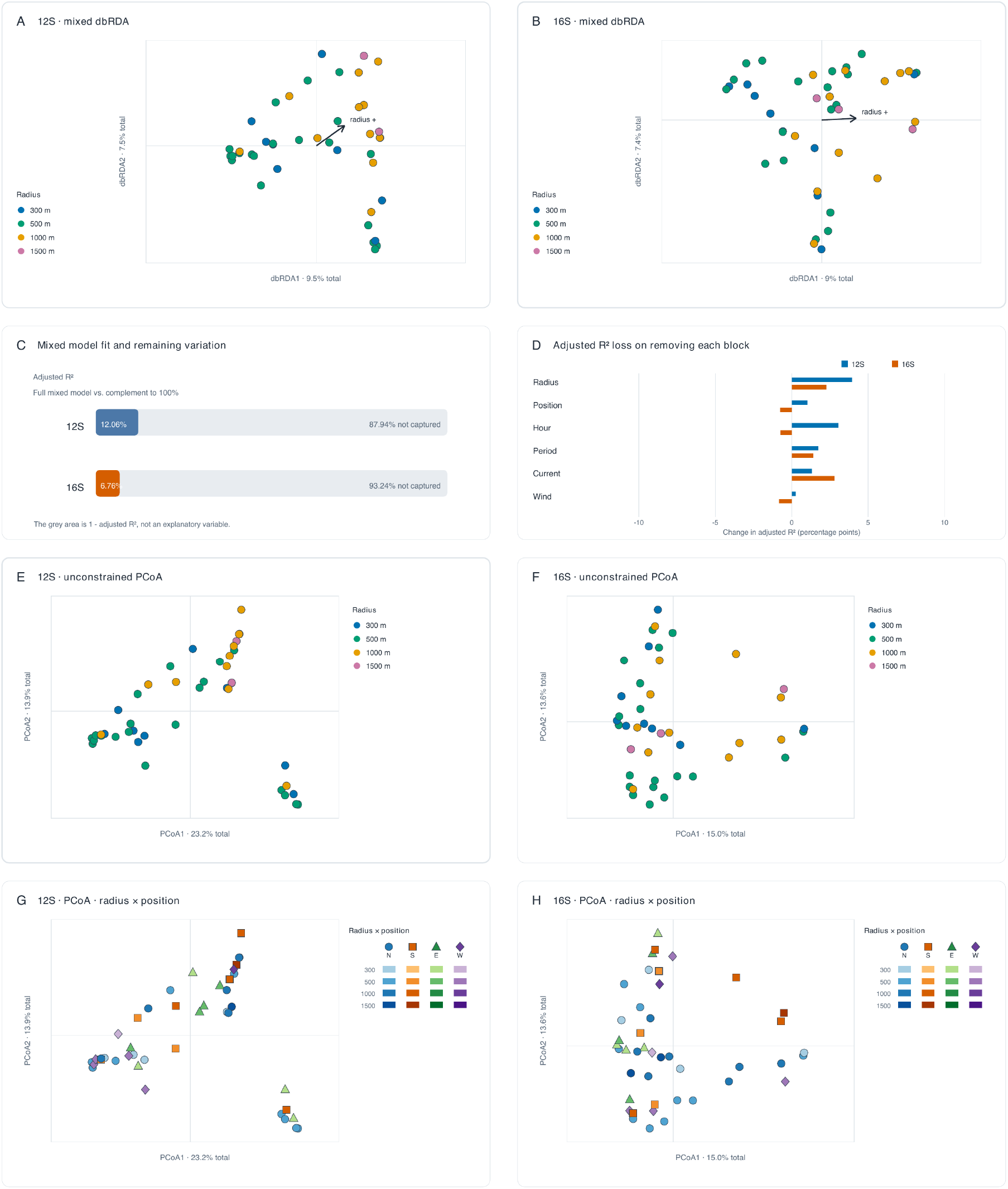
Local community composition, model fit and spatial grouping around the FPSO. **A–B**. Distance-based redundancy analysis (dbRDA) for 12S and 16S, respectively, constrained by planned radius, position relative to the platform, sampling hour, period of day, and current and wind direction and speed. Each point is a sample/sample; blue, green, orange and pink denote planned radii of 300, 500, 1,000 and 1,500 m. The arrow indicates the direction of association with increasing radius. **C**. Full-model adjusted R^2^ and its complement to 100% (grey; 87.94% and 93.24%). The grey portion is variation not captured by the model, not a fitted residual predictor. **D**. Change in adjusted R^2^ after removing each predictor block while retaining all other blocks and the same samples. Blue and orange bars denote 12S and 16S. Positive values indicate loss of adjusted fit after removal; negative values indicate improved adjusted fit. **E–F**. Unconstrained principal coordinates analysis (PCoA) of Euclidean distances among the same Hellinger-transformed profiles, coloured by radius as in A–B. **G–H**, the identical PCoA coordinates, now coded jointly by radius and directional sector.

We also tested assignment associations with near-platform (300–500 m) versus farther (1,000–1,500 m) samples using the in-house Python script `indval_local_43.py`, documented with NumPy 2.0.2. Specificity was defined as mean within-sample relative abundance in a group divided by the sum of group means; fidelity was defined as the proportion of that group’s samples with a nonzero detection. IndVal was 100× specificity × fidelity, and the test statistic was the maximum across the two groups. All 81 detected 12S assignments and 61 detected 16S assignments in the respective 43-sample cohorts were tested.

Three null schemes were compared: unrestricted sample-label permutation (199,999 random permutations; seed 42), exact point-label enumeration retaining 13 near and eight far points (203,490 assignments), and exact point-label enumeration within directional sectors (2,016 assignments). The point-level schemes kept repeated samples together; complete enumeration included the observed arrangement. Benjamini– Hochberg correction at 5% was applied separately within each marker over its full tested assignment family.

### Technical reproducibility and short-term temporal variability

#### Technical, within-point and between-point dissimilarity

We compared Hellinger dissimilarities at three levels: paired manual and Opentrons preparations from the same source DNA, repeated biological samples at the same point over approximately 1–2 h, and samples from different points. Technical comparisons included 11 source–replicate pairs for 12S and nine for 16S after read-depth filtering. Biological pairwise comparisons were restricted to 14 reoccupied points, comprising 39 samples for 12S and 38 for 16S. These yielded 36 and 34 within-point dyads and 705 and 669 between-point dyads, respectively; singleton-point samples did not enter these dyad distributions.

The in-house Python script `community_dissimilarity_ permutation_tests.py` tested two one-sided planned contrasts per marker using differences in median dissimilarity. For technical versus temporal variation, source identity was reassigned among qualifying biological samples at the same point while holding the technical profiles and temporal median fixed. All combinations of eligible source assignments were enumerated: 177,147 for 12S and 13,122 for 16S. For temporal versus spatial variation, point labels were permuted 9,999 times among the biological samples, preserving the observed number of samples per point. The test statistic was the between-point median minus the within-point median; Monte Carlo p-values included the observed arrangement through the correction (b + 1)/(B + 1). Holm adjustment was applied across the four marker-by-contrast tests. Randomized point-label tests used seed 42. The permutation units were source identity or sample labels, not rows of pairwise distances, because dyads sharing a sample are dependent.

Supplementary Figure S1B–C reused one PCoA per marker (12S: 46 biological and 11 technical profiles; 16S: 45 and nine), computed by eigen decomposition of the double-centred squared-distance matrix; axis percentages used total positive inertia. Nearest neighbours were determined in full Hellinger space, excluding self-matches. Bidirectional source–replicate comparisons used all qualifying local profiles; same-point comparisons queried biological samples from reoccupied points against all qualifying biological profiles, including singleton points.

An additional processing-group comparison used vegan’s `adonis2` in the in-house R script `13-replicas.R`, with Hellinger-transformed local profiles as the response, manual versus Opentrons preparation as the grouping factor, Euclidean distances and 999 unrestricted permutations (seed 42). This was a sample-level comparison of processing groups, not a paired or point-restricted permutation test. Technical replicates assessed the laboratory stages repeated in the protocol comparison, not independent field sample or DNA extraction.

#### Temporal turnover and nestedness

We partitioned Jaccard dissimilarity into turnover and nestedness-resultant components with the in-house R script `17-turnover-nestedness.R`, implementing the Baselga pairwise formulation directly. For each pair, a was the number of shared taxa and b and c the numbers unique to each sample. Total dissimilarity was (b + c)/(a + b + c), turnover was 2 min(b,c)/[a + 2 min(b,c)], and nestedness was the difference between total dissimilarity and turnover. The analysis used the same read-filtered biological samples and within- and between-point dyads described above, initially defining presence as at least one read.

Components were averaged over dyads within each comparison class. Percentages reported in Table 2 were calculated as the mean component divided by mean total Jaccard dissimilarity, not the mean of pairwise component percentages. Sensitivity analyses repeated the decomposition at presence thresholds of 5, 10 and 20 reads. Mean within-point components were also retained by geographic point. The decomposition described taxon replacement and subset patterns; it did not identify their biological or transport mechanisms. Dyads were not treated as independent field replicates.

**Table 2.**
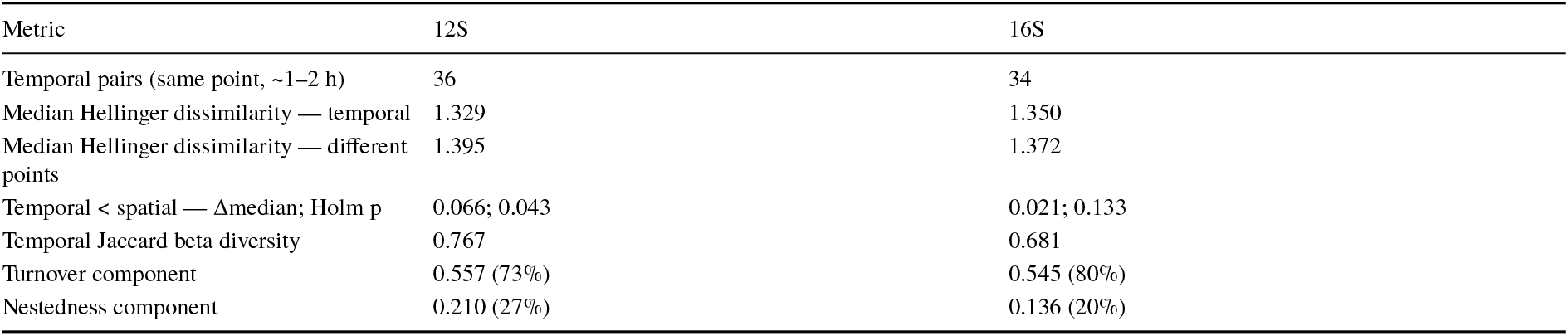
Short-term temporal variability and beta-diversity decomposition.

### Regional community differentiation and indicator assignments

#### Regional constrained ordinations

Regional differentiation was evaluated using the in-house R script `08-dbrda.R` and vegan’s `dbrda` function. Five representations were fitted: 12S taxa (63 biological samples), 16S taxa (62), combined taxa (64), 12S OTUs (63) and 16S OTUs (62). Models used Hellinger-transformed profiles and Euclidean distances, with distance from the FPSO in kilometres and sampling date as a numeric day. The combined model used the separately normalized, summed marker profiles described in Section 2.5; its read-depth filter came from summed raw taxonomic counts, not the normalized matrix. Distance was calculated from geographic coordinates using an equirectangular approximation with Earth radius 6,371 km and FPSO reference coordinates 25.475852° S, 43.938755° W.

Global model tests used 999 unrestricted permutations with seed 42. Adjusted R^2^ and axis percentages were calculated from the fitted models; percentages referred to total inertia. Supplementary Figure S2 was assembled through the in-house script `figure_s3_regional.py`. Taxonomic models were selectively refitted after curation; OTU fits were retained. Updated taxonomic scores and annotations were unchanged at the displayed precision. Regional models were distinct from the local complete-case models. Their unrestricted tests did not keep repeated samples together, and locality, distance and campaign date were confounded; they therefore did not isolate independent causal effects of these predictors.

### Exploratory regional indicator-value analysis

The in-house R script `10b-indicadores.R` implemented IndVal directly using the specificity and fidelity definitions in Section 2.7. Groups were coast (Urca plus Ilhas Cagarras; eight samples), transition (transit; nine) and offshore (FPSO; 46 samples for 12S and 45 for 16S). The statistic was the maximum IndVal across the three groups. Its maximizing group was re-selected in each of 999 unrestricted sample-label permutations, and p-values used the correction (b + 1)/(999 + 1). The script used seed 42.

Associations were screened at nominal, unadjusted p < 0.05; no multiple-testing correction or point-level permutation restriction was applied to the regional screen reported in Fig. 4C. These associations remained exploratory. The figure displays a selected subset of nominal associations, rather than all assignments passing the screen. The in-house Python script `regional_indval_components.py` recovered specificity, fidelity and IndVal for each displayed entry in all three groups (Supplementary Table S2), retaining the original maximum-IndVal p-value. Components for nonselected groups were descriptive comparisons, not separate significance tests.

**Fig. 4.**
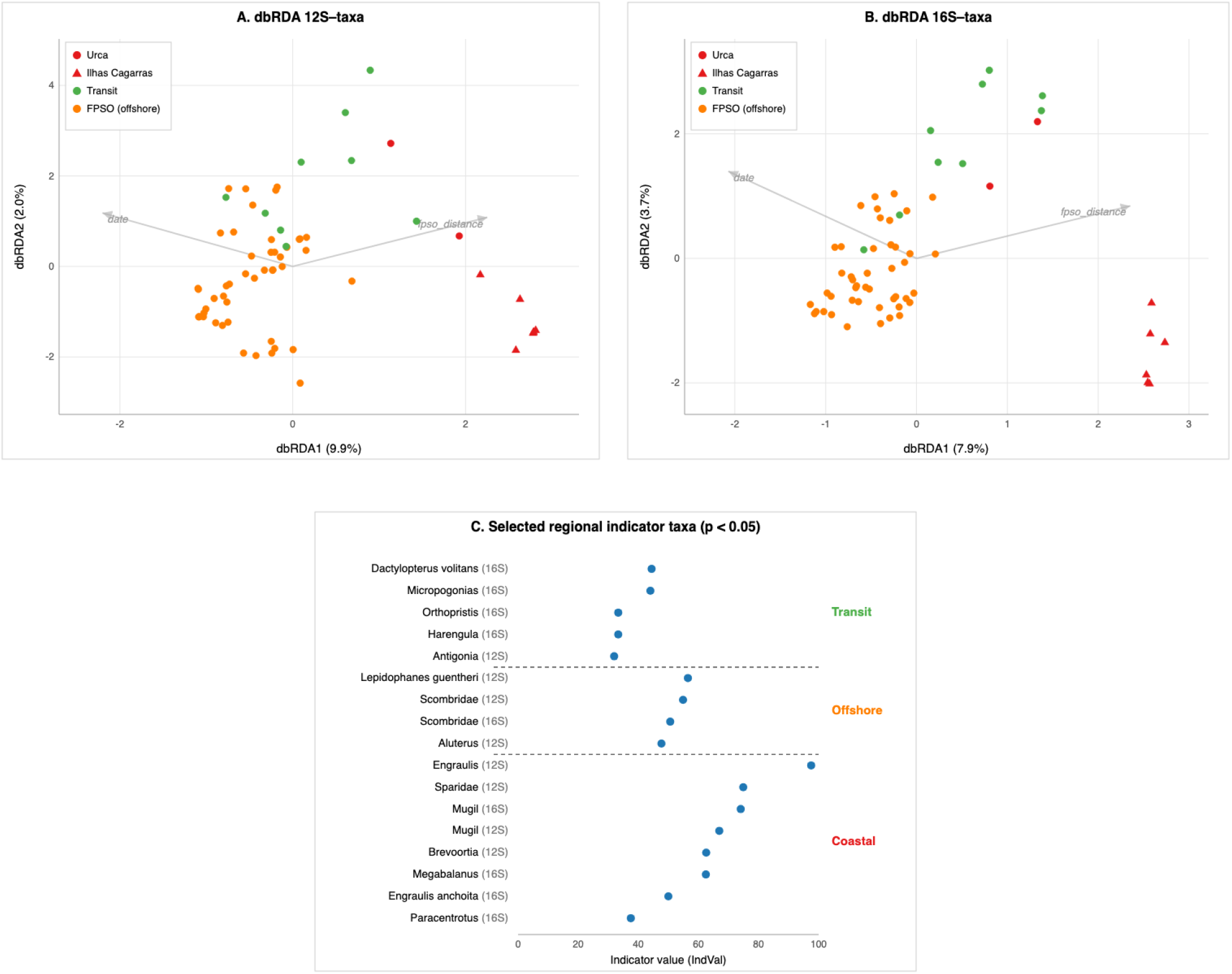
Regional community differentiation and exploratory indicator taxa. Regional analyses cover the coastal-to-offshore campaign from Urca and Ilhas Cagarras through transit locations to the floating production, storage and offloading unit (FPSO). Technical replicates are excluded. **A–B**, distance-based redundancy analysis (dbRDA) of Hellinger-transformed taxonomic read profiles for 12SMiFish (63 biological samples) and 16S-MarVer3 (62 samples), retaining libraries with at least 1,000 assigned reads. Models include distance from the FPSO and sampling date. Points represent samples. Red identifies the two coastal localities, with circles for Urca and triangles for Ilhas Cagarras; green circles identify transit samples and orange circles the offshore FPSO area. The vectors labelled “date” and “fpso_distance” indicate sampling date and distance from the FPSO, respectively. Axis labels report percentages of total inertia: 9.9% and 2.0% for the first two 12S axes, and 7.9% and 3.7% for 16S; these are not percentages of constrained inertia alone. **C**, selected taxonomic assignments from a separate regional indicator-value analysis (IndVal), grouped as Transit, Offshore (FPSO) and Coastal (Urca plus Ilhas Cagarras). Each dot gives IndVal on a 0–100 scale, and the marker is identified after the taxon name. IndVal is 100 × specificity × fidelity: specificity is the group’s mean relative read abundance divided by the sum of group means, and fidelity is the proportion of samples in that group with a nonzero detection. The displayed assignments passed an unadjusted permutation p < 0.05 threshold (999 permutations); no correction for testing multiple assignments was applied, so they remain exploratory associations. Dashed separators distinguish the three groups. Group-specific IndVal and its specificity and fidelity components for every displayed entry are provided in Supplementary Table S2. The regional dbRDA associations do not isolate an independent distance effect because locality, distance and campaign date are confounded.

## Results

The final dataset comprised 212 12S OTUs and 241 16S OTUs, represented by 106,605,095 and 26,934,807 reads, respectively, across 75 libraries (Supplementary Data 5a–b). Taxonomic assignments were obtained for 205 and 219 OTUs, respectively, whereas seven 12S and 22 16S OTUs remained unassigned.

The taxonomic matrices contained 103 taxa for 12S and 131 for 16S, spanning species-level to higher-rank assignments. Thirty-five taxa were shared between markers, yielding 199 distinct assigned taxa overall (Supplementary Data 4a–b).

### Local taxonomic diversity and sampling completeness

The local survey detected 34 species with 12S and 28 with 16S across 47 biological samples from 21 geographic sampling points (Fig. 2A–B; Supplementary Data 4a–b). For 12S, the observed inventory represented 74.3% of the incidence-based Chao2 richness estimate of 45.7 species; the corresponding 16S values were 65.4% and 42.8 species. Both sample-based curves showed diminishing gains with increasing effort, providing a quantitative reference for extending the inventory. Doubling the effort to 94 samples was projected to increase detected richness to 41.5 species for 12S and 35.7 for 16S (Supplementary Figure S3A). The continued increase in the curves is consistent with the large fraction of species and OTUs detected in only one or two samples (singletons and doubletons; Supplementary Figure S3C). Together, these low-incidence units accounted for 18 of 34 species and 97 of 142 OTUs for 12S, and 15 of 28 species and 98 of 135 OTUs for 16S. Removing singletons flattened the extrapolated point estimates (Supplementary Figure S3B), illustrating their contribution to estimated unseen richness. The survey thus established an initial species inventory while identifying the additional detections expected from greater sampling effort.

The combined profiles highlighted recurrent detections of both fishes and planktonic taxa (Fig. 2C). Ray-finned fishes (Actinopteri) accounted for 42 of the 50 displayed taxa, with the remaining eight spanning Hydrozoa, Thaliacea, Hexanauplia, Mammalia, and Asteroidea. This predominance extended beyond the displayed top 50: Actinopteri accounted for 106 of 129 detected taxa (82.2%) in the complete local taxonomic matrices and 74,658,784 of 77,029,888 taxonomically assigned reads (96.92%; Supplementary Table S1). Marker-specific proportions were 78 of 85 taxa (91.8%) and 99.97% of assigned reads for 12S, versus 47 of 65 taxa (72.3%) and 84.66% of assigned reads for 16S. A plausible contribution to this pattern is the choice of assays: MiFish was developed for fish metabarcoding [18], whereas MarVer3 was designed for marine vertebrates, including marine mammals [19].

The family-level Scombridae assignment occurred in 35 of 47 samples (74.5%), and the lanternfish *Lepidophanes guentheri* was the most prevalent displayed fish species, occurring in 31 (66.0%). Other recurrent fish assignments included *Aluterus* (22 samples, 46.8%), the gray triggerfish *Balistes capriscus* (16, 34.0%), and *Diaphus perspicillatus*, another member of the lanternfish family Myctophidae (14, 29.8%). Among nonfish assignments, the hydrozoans *Aglaura hemistoma* and *Liriope tetraphylla* occurred in 39 (83.0%) and 37 (78.7%) samples, respectively, while the pelagic tunicate *Doliolum nationalis* also occurred in 37 (78.7%). The profiles also contained an assignment to Delphinidae, the oceanic dolphin family, in seven samples (14.9%), extending the detected signal to marine mammals. This assignment was resolved only to family level and therefore did not identify a particular dolphin species. Together, these detections illustrate the taxonomic breadth of the eDNA inventory, from hydrozoans and pelagic tunicates to ray-finned fishes and marine mammals.

### Local composition shows no strong and consistent radial gradient around the FPSO

We fitted local dbRDA models to 43 biological samples per marker using planned radius (300–1,500 m), relative position, sampling hour, period of day, and current and wind direction and speed. The full models had modest adjusted R^2^ values: 0.1206 for 12S and 0.0676 for 16S (Table 3). Removing each predictor block in turn, while retaining the others, identified radius as a consistently contributing predictor: its removal reduced adjusted R^2^ by 3.95 percentage points for 12S and 2.27 for 16S, the largest and second-largest losses in the respective markers (Fig. 3D). Nevertheless, losses were small across all blocks, and most variation remained uncaptured: 87.94% for 12S and 93.24% for 16S, expressed as the complement of adjusted R^2^ (Fig. 3C). To test whether current and wind improved the model fit beyond the sampling design, we also compared models with and without these covariates on the same samples. Adjusted R^2^ increased from 0.101 to 0.121 for 12S and from 0.031 to 0.068 for 16S, but neither improvement was statistically significant (nested permutation tests: p = 0.235 and 0.076, respectively).

**Table 3.** Full local dbRDA models using 43 complete-case biological samples per marker.

| Marker | n | Pseudo-F | Permutation p | Adjusted R<sup>2</sup> | dbRDA1 (% total inertia) | dbRDA2 (% total inertia) |
| --- | --- | --- | --- | --- | --- | --- |
| 12S | 43 | 1.443 | 0.003 | 0.1206 | 9.5 | 7.5 |
| 16S | 43 | 1.234 | 0.026 | 0.0676 | 9.0 | 7.4 |

Despite the consistent contribution of radius to model fit, the first two dbRDA axes showed no clear separation of samples coloured by radius (Fig. 3A–B). We therefore complemented the constrained ordinations with PCoA of the same Hellinger-transformed community profiles. Unlike dbRDA, PCoA summarizes overall compositional dissimilarities without using the measured predictors to constrain its axes, allowing us to examine whether radial grouping was apparent in the dominant patterns of community variation. We first coloured samples by radius (Fig. 3E–F), then displayed the same coordinates using both radius and relative position around the FPSO (Fig. 3G–H). Neither display revealed clear, consistent separation among groups in the first two axes. Thus, the contribution of radius to the fitted models did not translate into distinct radial groups in the displayed ordinations.

A separate exploratory indicator-value analysis (IndVal) tested whether taxonomic assignments were associated with near-platform samples (300–500 m) or farther samples (1,000–1,500 m). Using the same 43 samples per marker as the local dbRDA models, we tested 81 taxonomic assignments for 12S and 61 for 16S. No assignment remained significant after Benjamini–Hochberg correction at 5% within each marker. This result held for unrestricted sample-level permutations and for point-level permutations that kept repeated samples together, both with and without stratification by quadrant. Thus, this analysis did not identify taxonomic indicators of near- or far-platform samples after correction for multiple testing.

#### Short-term compositional variation exceeds technical variation and approaches between-point differences

Water samples made at the same point over approximately 1– 2 h were almost as dissimilar as samples from different points (Supplementary Figure S1A; Table 2). Median Hellinger dissimilarities were 1.329 versus 1.395 for 12S and 1.350 versus 1.372 for 16S; the small within-point similarity advantage was significant only for 12S (Holm-adjusted p = 0.043 and 0.133, respectively). In contrast, technical replicates closely matched their source DNA, with median dissimilarities of 0.015 for 12S and 0.037 for 16S, both lower than the temporal comparisons (Holm-adjusted p = 0.011 and 0.019; Supplementary Figure S1A–B). Thus, variability in the laboratory-processing steps tested through technical replicates was unlikely to account for the larger differences between repeated field samples.

Nearest-neighbour relationships reinforced this contrast. Every technical replicate had its source DNA profile as its nearest neighbour in the full Hellinger distance matrix (11/11 for 12S and 9/9 for 16S; Supplementary Figure S1B). Among repeated field samples, however, only 5 of 39 for 12S and 3 of 38 for 16S had their nearest biological neighbour at the same geographic point (Supplementary Figure S1C). Samples from the same point therefore rarely shared the most similar community profiles, i.e. generally were similar to samples from other points.

Temporal differences were dominated by turnover of detected taxa rather than nestedness, in which one sample contains a subset of the taxa in another. Turnover accounted for approximately 73% of mean temporal Jaccard dissimilarity in 12S and 80% in 16S, compared with 27% and 20% for nestedness (Table 2). Turnover remained dominant when the detection threshold increased from one read to 5, 10 or 20 reads. These results identify taxon replacement as the main compositional pattern.

### eDNA recovers a consistent regional environmental contrast

Unlike the overlapping radial groups within the local FPSO sampling area, the regional dbRDA plots showed grouping by geographic origin, most clearly distinguishing coastal from offshore samples, with some overlap involving transit samples (Fig. 4A–B; Supplementary Figure S2). This contrast is consistent with the broader coastal-to-offshore sampling extent encompassing compositional differences that were not clearly resolved across the 300–1,500 m local design. Regional models including distance from the FPSO and sampling date were significant for all five representations — 12S taxa, 16S taxa, combined taxa, 12S OTUs and 16S OTUs (all p = 0.001) — with adjusted R^2^ values of 0.089, 0.087, 0.098, 0.095 and 0.086, respectively (Table 4). The pattern therefore supports a scale-dependent interpretation of the observed differentiation. We then used an exploratory indicator-value analysis (IndVal) to identify taxonomic assignments associated with each regional group: coast (Urca and Ilhas Cagarras), transition (transit) and offshore (FPSO). At the nominal, unadjusted p < 0.05 threshold, 12S yielded 14 coastal, two transitional and four offshore assignments, whereas 16S yielded 13 coastalten transitional and one offshore assignment. Fig. 4C shows selected associations, and Supplementary Table S2 compares their IndVal, specificity, fidelity and original p-values across the three groups.

**Table 4.** Regional compositional models across the coastal-to-offshore campaign.

| Marker / representation | n | Effect / model | p | Adjusted R<sup>2</sup> / effect |
| --- | --- | --- | --- | --- |
| 12S taxa | 63 | distance + date | 0.001 | 0.089 |
| 16S taxa | 62 | distance + date | 0.001 | 0.087 |
| Combined taxa | 64 | distance + date | 0.001 | 0.098 |
| 12S OTUs | 63 | distance + date | 0.001 | 0.095 |
| 16S OTUs | 62 | distance + date | 0.001 | 0.086 |

The coastal associations included *Mugil* (mullets) in both markers, detected in 87.5% of coastal samples for 12S and 75.0% for 16S. This association is ecologically consistent with the use of coastal waters and estuaries by members of the genus, such as *M. liza*, without resolving the genus-level detection to that species [24]. Other coastal assignments included *Engraulis* (anchovies) in 12S and *Megabalanus* (acorn barnacles) in 16S. These signals are compatible with coastal pelagic fishes and hard-substrate-associated invertebrates, respectively, although neither group is uniquely coastal: *Engraulis* includes species that range offshore, and *Megabalanus* includes settlers on artificial surfaces [25, 26]. In the offshore group, *Lepidophanes guentheri*, an oceanic lanternfish (family Myctophidae), was detected by 12S in 26 of 46 samples (56.5%; Supplementary Table S2). Its association with this group is consistent with its oceanic habitat [27].

## Discussion

This study demonstrates the potential of autonomous sampling coupled with eDNA metabarcoding for offshore biodiversity monitoring. The local survey comprised 47 biological samples at 21 geographic points. Within the subset of repeatedly sampled points, profiles differed substantially over approximately 1–2 h, while the local spatial analyses identified weak radial differentiation across the 300–1,500 m sampling range. Autonomous sampling offers a means of expanding repeated observations to characterize this short-term variability.

The local composition was associated with the measured predictors, but the fitted models captured only a small fraction of its variation. Radius contributed to model fit without producing distinct radial groups in the ordinations (Fig. 3; Table 3). Likewise, no taxonomic assignment remained associated with near- or far-platform samples after correction for multiple testing, including analyses that kept repeated samples from the same point together. Together, these results describe a weak spatial pattern distributed across overlapping profiles, rather than distinct assemblages organized by distance from the FPSO. Adding current and wind direction and speed did not significantly improve the sampling-design models.

Samples collected at the same point approximately 1–2 h apart were almost as dissimilar as samples from different points, whereas technical profiles closely matched their source DNA (Table 2). A small within-point similarity advantage remained for 12S, but was not statistically detected for 16S. Returning to the same coordinate therefore did not reliably recover a similar molecular profile. The predominance of turnover, which persisted at higher read-presence thresholds, indicates that repeated sampling captured different sets of taxa rather than mainly adding or losing subsets of the same assemblage. Temporal replication thus helps characterize the range of detections at a location instead of simply reproducing an initial observation. The agreement between technical profiles supports this interpretation for the post-extraction steps tested, while independent sampling and consistent processing remain important for accounting for detection variability [28, 29].

The turnover observed between repeated samples could partly reflect changes in nearbyDNAsources. Proximity to organisms can sustain detectable signals, while dilution and settling attenuate material transported from more distant sources [30]. Organisms passing near a sampling point could therefore contribute DNA detected in one sample that subsequently disperses below detection. This balance between renewed local inputs and attenuation offers a plausible explanation for changing detections at a fixed location, without requiring a persistent radial gradient [30].

The two markers provide complementary views of the sample assemblage. Fish reads predominated in both, but the larger nonfish contribution to 16S profiles shows why pooling the markers can obscure differences in taxonomic coverage (Supplementary Table S1). This pattern is consistent with the targeting of fishes by MiFish and marine vertebrates by MarVer primers [18, 19]. Broader coverage may also reveal environmental gradients expressed by other community components: vertical differentiation involving dinoflagellates and copepods can be more pronounced in broad-eukaryote 18S profiles than in fish-focused MiFish profiles [7]. Adding 18S or COI assays could therefore complement the vertebrate inventory with taxa whose spatial and temporal distributions differ [7, 15]. Whether this improves the detection of environmental change would depend on the patterns expressed by those additional components. Shotgun sequencing has already been performed on these samples, and its forthcoming analysis will assess whether broader molecular characterization reveals patterns not captured by the two amplicon assays.

A single three-day campaign established an initial inventory of 34 species with 12S and 28 with 16S. Although neither curve reached a plateau, both showed diminishing gains with increasing effort. Doubling the number of samples from 47 to 94 was projected to add approximately seven to eight species per marker under comparable sampling conditions (Supplementary Figure S3A). The campaign therefore recovered a substantial share of the estimated species inventory, while leaving scope for additional detections. More than half of the detected species occurred in only one or two samples. Such detections may reflect intermittent occurrence or imperfect detection rather than low organism abundance [29]. Extending observations across seasons can reveal taxa and community states poorly represented within a short campaign [2]. Repeated sampling would therefore help distinguish recurrent, seasonal and occasional detections, while additional taxa may continue to accumulate even as frequently detected components become better characterized.

Weak radial grouping coexisted with a regional compositional pattern across markers and both taxonomic and OTU-based analyses. The regional models were significant but captured only a modest fraction of overall variation, indicating a consistent contrast within otherwise heterogeneous profiles (Table 4). Coastal associations such as the genus-level *Mugil* assignment in both markers, and offshore associations such as *Lepidophanes guentheri* in 12S and Scombridae in 16S, illustrate its taxonomic expression. These indicator profiles remain exploratory. The regional inference concerns the campaign as sampled, with geographic and temporal variation jointly represented, rather than an isolated effect of distance.

The reference characterization established here applies to eDNA collected at 1 m depth during a three-day campaign, at sampling points 300–1,500 m from the FPSO. Extending monitoring closer to the structure and at different depths could test for patterns that additional samples within the same 300– 1,500 m radii may not capture. Platform-associated differences can be concentrated within approximately 50 m and be more pronounced in benthic assemblages [6], while fish and plankton profiles can also differ between depths [7].

Autonomous sampling provides a means of extending offshore observations across the time scales needed for biodiversity monitoring. The results support additional sampling effort, while short-term turnover and the need to characterize recurrence support distributing that effort through time. Marker complementarity further supports evaluating multiple community components. Together, these results favour frequent sampling sustained across campaigns, so that transient detections can be distinguished from recurrent patterns and persistent compositional changes.

## Conclusions

This pilot shows that eDNA metabarcoding, coupled with autonomous offshore sampling, can provide a technically reproducible and ecologically informative basis for biodiversity monitoring around offshore infrastructure. Rather than providing evidence of a platform effect, the study identifies the spatial and temporal scales of variability that must be incorporated into future monitoring designs. This distinction is critical for separating short-term ecological turnover from persistent change and for defining monitoring effort according to specific biodiversity endpoints.

## Supporting information

supplementary-data-manifest

taxonomic-curation-map-20261019

taxonomic-curation-log-20261019

supplementary-table-s2-indval-components.

supplementary-table-s1-class-composition

supplementary-data-sample-map

supplementary-data-guide

supplementary-data-crosscheck

supplementary-data-contaminant-exclusions

supplementary-data-5b-otus-16S

supplementary-data-5a-otus-12S

supplementary-data-4b-taxa-16S

supplementary-data-4a-taxa-12S

supplementary-data-3b-processing-16S

supplementary-data-3a-processing-12S

supplementary-data-2-pcr-thermocycling

supplementary-data-1-sampling

## Supplementary material

**Fig. S1.**
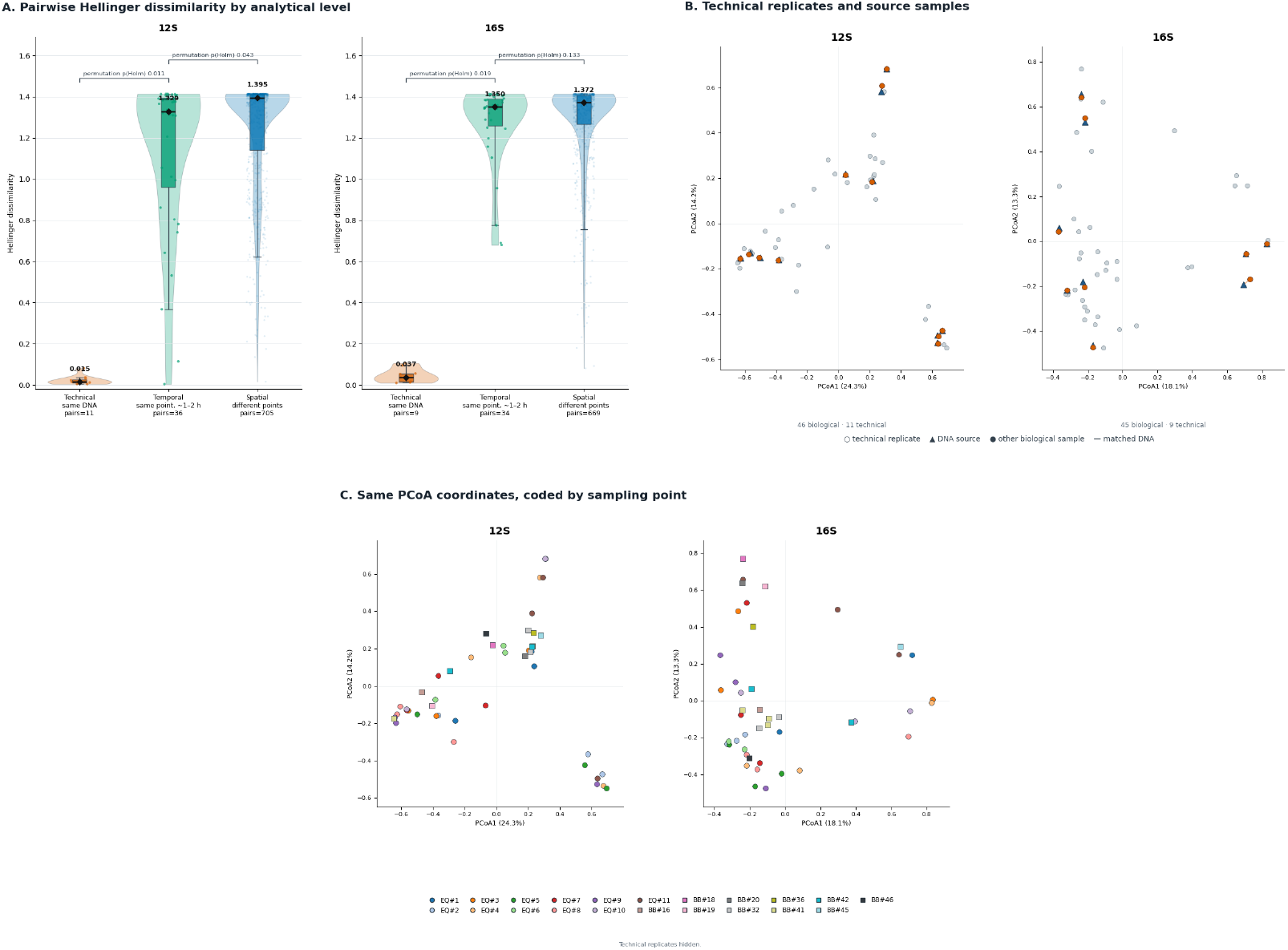
S1 Fig. Technical reproducibility compared with short-term biological variability around the FPSO. Analyses use 12S-MiFish and 16S-MarVer3 taxonomic profiles from the local sampling area, retaining profiles with at least 1,000 assigned reads. Hellinger dissimilarity is the Euclidean distance between square-root-transformed relative read profiles; smaller values indicate more similar composition. **A**, distributions of pairwise dissimilarities for technical replicates versus their source DNA (orange; 11 pairs for 12S and nine for 16S), repeated biological samples at the same point over approximately 1–2 h (green; 36 and 34 pairs), and samples at different points (blue; 705 and 669 pairs). The biological comparisons use 39 12S and 38 16S samples from 14 reoccupied points. Dots are individual pairwise distances with horizontal jitter; violins show their distributions, boxes span the interquartile range, black lines and adjacent numbers mark medians, and whiskers indicate the 5th and 95th percentiles. Brackets report one-sided permutation tests of lower technical than temporal dissimilarity and lower temporal than spatial dissimilarity, with Holm adjustment across the four marker-by-contrast tests. Source identities were reassigned within sampling points for the technical–temporal contrast; point labels were permuted while preserving point sizes for the temporal–spatial contrast. Pairwise distances sharing a sample are not independent replicates. **B**, principal coordinates analysis (PCoA) of the full local Hellinger distance matrix for each marker: 46 biological plus 11 technical profiles for 12S, and 45 biological plus nine technical profiles for 16S. Orange circles identify technical replicates, blue triangles identify their source DNA profiles, grey circles are other biological samples, and lines connect matched source–replicate pairs. **C**, the same PCoA coordinates with technical replicates hidden and biological samples identified by sampling-point colour and symbol; circles denote EQ point IDs and squares denote BB point IDs. Matching IDs indicate the same geographic point. Axis percentages in B–C refer to total ordination inertia. Nearest neighbours were determined in the full Hellinger distance space, not from the two plotted axes. Technical comparisons allow all qualifying local profiles as neighbours; same-point biological counts allow only biological profiles and evaluate samples from reoccupied points. The technical-batch p-values come from permutational multivariate analysis of variance (PERMANOVA). Technical replicates assess the laboratory-processing comparison, not independent field replication.

### Complementary technical-reproducibility results (Supplementary Figure S1B)

Every technical replicate had its own source sample as its nearest neighbor in the full Hellinger distance matrix (11/11 for 12S and 9/9 for 16S). In the reverse comparison, the technical replicate was the nearest neighbor of its source in 9/11 and 9/9 cases, respectively. The FPSOrestricted PERMANOVA comparing technical-replicate and biological profiles detected no processing-group difference (R^2^ ≈ 0.009 for both markers; p = 0.909 for 12S and p = 0.959 for 16S).

**Fig. S2.**
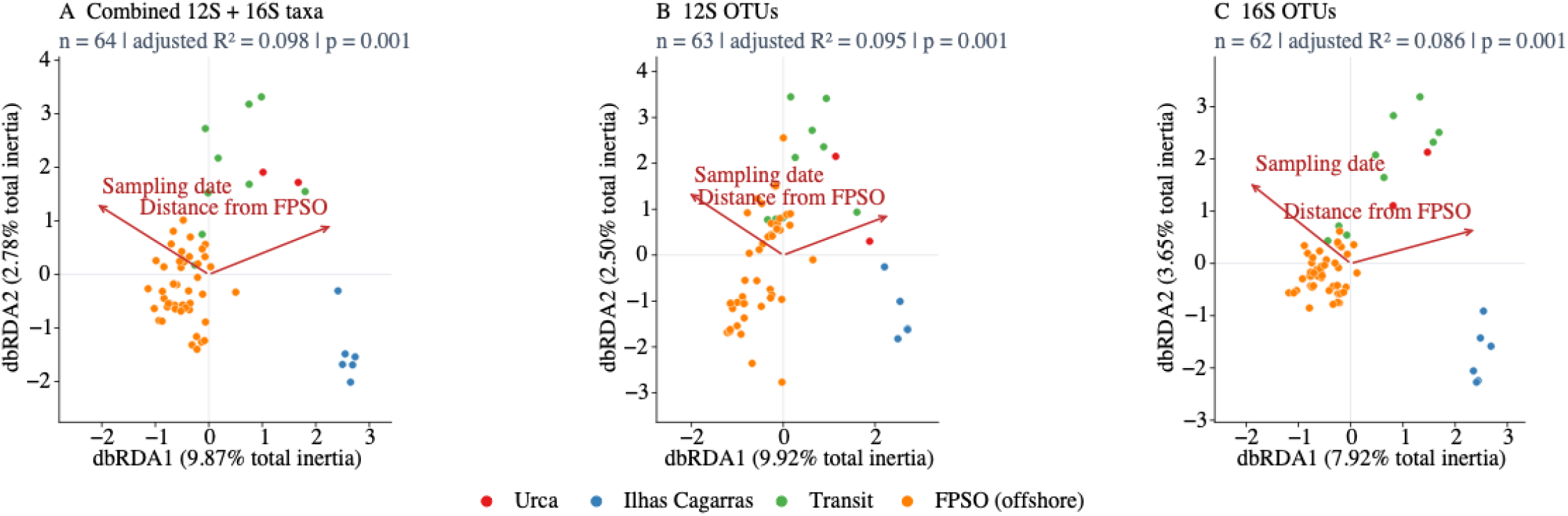
S2 Fig. Regional dbRDA of combined taxonomic profiles and marker-specific OTUs. The panels complement the separate 12S and 16S taxonomic ordinations in Fig. 4. **A**, combined 12S-MiFish and 16S-MarVer3 taxonomic profiles (64 biological samples); **B**, 12S operational taxonomic units (OTUs; 63 samples); **C**, 16S OTUs (62 samples). The regional campaign comprises Urca, Ilhas Cagarras, transit locations and the floating production, storage and offloading unit (FPSO). Each point represents one biological sample; technical replicates are excluded. Red, blue, green and orange identify Urca, Ilhas Cagarras, transit and the offshore FPSO area, respectively, using the same regional colours as Fig. 4. Profiles were Hellinger transformed (square roots of within-sample relative values), and distance-based redundancy analysis (dbRDA) used Euclidean distances with distance from the FPSO and sampling date as predictors. The combined taxonomic input sums the separately normalized marker profiles, giving equal weight to available marker profiles before Hellinger transformation; OTU inputs use marker-specific read counts. Libraries were retained at a minimum of 1,000 reads in the corresponding input; the combined representation uses the summed taxonomic read count for this filter, not separate 1,000-read requirements for each marker. Red arrows show the saved biplot vectors for increasing distance and date. Site and vector scores use vegan scaling 2; vectors are multiplied by 2.5 for display, as in Fig. 4, so their lengths should not be read as effect sizes in site-score units. Axis percentages refer to total inertia, not only constrained inertia. Panel annotations give the sample count, adjusted R^2^ and full-model permutation p-value (999 permutations). Each panel has its own coordinate limits, with equal horizontal and vertical units; absolute coordinates are not comparable across independently fitted representations. The combined taxonomic model was checked against the selectively refitted curated model; scores and annotations were unchanged at the displayed precision. The OTU panels retain the original fitted models and scores. Locality, distance and campaign date are confounded, so the arrows do not identify independent causal effects.

**Fig. S3.**
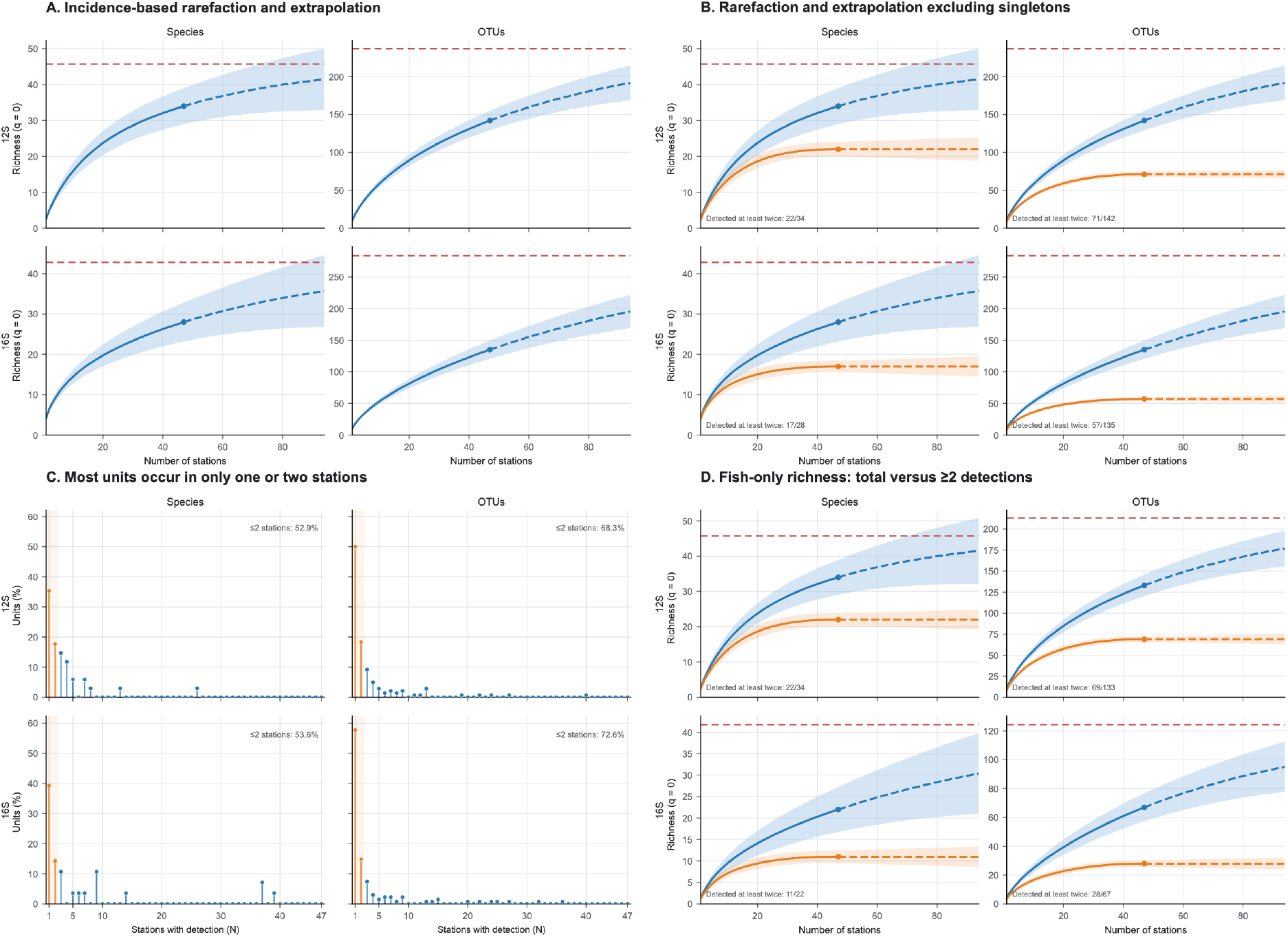
S3 Fig. Monitoring endpoint determines sampling sufficiency. Analyses use 47 biological water samples from 21 geographic points around the floating production, storage and offloading unit (FPSO), excluding additional technical replicates. The historical axis label “stations” denotes biological samples, not 47 distinct geographic points. Within each panel, rows show 12S-MiFish above 16S-MarVer3 and columns show species on the left and operational taxonomic units (OTUs) on the right. Species and OTUs are separate observation units; multiple OTUs assigned to one species remain distinct. Incidence records whether a unit was detected in a sample. **A**, incidence-based richness (Hill number q = 0) rarefaction through the observed 47 samples and extrapolation to 94. Solid blue curves denote rarefaction, dashed blue segments denote extrapolation, points mark observed richness, and horizontal dashed red lines mark Chao2 richness estimates. Shading represents 95% parametric-bootstrap confidence intervals based on 200 replicates. **B**, the same all-unit curves in blue compared with orange curves restricted to units detected in at least two of the complete set of 47 samples. Subset membership is fixed before rarefaction and all sample rows are retained. The ≥2-detection subsets contain 22 of 34 species and 71 of 142 OTUs for 12S, and 17 of 28 species and 57 of 135 OTUs for 16S. In B and D, solid segments denote rarefaction through 47 samples and dashed segments denote extrapolation beyond 47; circles mark richness observed at 47 samples for each blue or orange series, and matching blue or orange shading represents 95% parametric-bootstrap confidence intervals. Horizontal dashed red lines give the Chao2 estimate for all units in B and all fish units in D. Blue curves show all units in B and all fish units in D; orange curves show the corresponding subsets detected in at least two samples. B and D use the same incidence-based richness estimator as A. **C**, distribution of detection frequencies: the horizontal axis gives the number of samples containing a unit and the vertical axis the percentage of observed units with that frequency. Orange marks highlight singletons and doubletons (units detected in one or two samples); annotations give their combined percentage. Doubletons belong both to the highlighted group in C and to the retained orange subset in B. **D**, the comparison in B restricted to assignments with lineages in Actinopteri or Chondrichthyes. This is a fish filter, not a marine-only filter; OTUs without an established fish assignment are excluded. The ≥2-detection/total-observed counts are 22/34 species and 69/133 OTUs for 12S, and 11/22 species and 28/67 OTUs for 16S. Axes in D use fish-specific richness limits. Removing singletons forces estimated unseen richness under Chao2 to zero in the filtered input; the resulting orange plateau is therefore conditional on this retrospective filter, not evidence of a complete inventory or a prediction of future repeated detections. Intervals treat subset membership as fixed and do not incorporate selection uncertainty, taxonomic-assignment uncertainty or dependence among repeated field samples.

**Supplementary Table S1.** Complete local taxonomic composition by class. All 47 local biological samples are included, retaining standard-protocol profiles and excluding additional Opentrons profiles; no top-50 selection or multivariate read-depth exclusion is applied. Taxa are distinct detected curated taxonomic assignment IDs at their retained ranks, including assignments above species level, not species-richness estimates. Each ID is assigned to its recorded class; class-unresolved assignments remain explicit. Combined counts are the union of IDs across markers, not the sum of marker-specific counts. Reads are the unnormalized counts retained in the taxonomic matrices, summed across samples; percentages use all taxonomically assigned reads for the corresponding marker or pooled dataset, including class-unresolved reads. Pooled read proportions weight markers by their read totals and therefore differ from the equal-marker weighting in Fig. 2C; neither is organism abundance.

| Class | 12S taxa | 12S reads | 12S reads (%) | 16S taxa | 16S reads | 16S reads (%) | Combined taxa | Combined reads | Combined reads (%) |
| --- | --- | --- | --- | --- | --- | --- | --- | --- | --- |
| Actinopteri | 78 | 61,676,407 | 99.971 | 47 | 12,982,377 | 84.656 | 106 | 74,658,784 | 96.922 |
| Hydrozoa | 0 | 0 | 0.000 | 4 | 588,503 | 3.838 | 4 | 588,503 | 0.764 |
| Asteroidea | 1 | 13,771 | 0.022 | 2 | 34,920 | 0.228 | 3 | 48,691 | 0.063 |
| Hexanauplia | 0 | 0 | 0.000 | 3 | 123,262 | 0.804 | 3 | 123,262 | 0.160 |
| Holothuroidea | 0 | 0 | 0.000 | 2 | 372,263 | 2.427 | 2 | 372,263 | 0.483 |
| Mammalia | 2 | 4,209 | 0.007 | 1 | 100,105 | 0.653 | 2 | 104,314 | 0.135 |
| Thaliacea | 0 | 0 | 0.000 | 2 | 1,132,865 | 7.387 | 2 | 1,132,865 | 1.471 |
| Bivalvia | 0 | 0 | 0.000 | 1 | 7 | <0.001 | 1 | 7 | <0.001 |
| Cephalopoda | 0 | 0 | 0.000 | 1 | 139 | <0.001 | 1 | 139 | <0.001 |
| Magnoliopsida | 1 | 3 | <0.001 | 0 | 0 | 0.000 | 1 | 3 | <0.001 |
| Class unresolved | 3 | 38 | <0.001 | 2 | 1,019 | 0.007 | 4 | 1,057 | 0.001 |
| **Total** | **85** | **61,694,428** | **100.000** | **65** | **15,335,460** | **100.000** | **129** | **77,029,888** | **100.000** |

The corresponding OTU matrices contain 61,694,431 reads for 12S and 15,721,904 for 16S. The differences from the taxonomic-matrix totals are 3 unassigned 12S reads and, for 16S, 383,770 unassigned reads plus 2,674 reads assigned to Mammalia only in the OTU-assignment snapshot but absent from the taxonomic matrices. This documented source discrepancy is not silently merged into the table: totals describe the complete taxonomic matrices used for the profiles, not a reconstruction from every OTU assignment. Very small nonzero percentages are shown as <0.001 rather than zero. Taxonomic class names follow the supplied assignment lineages (Actinopteri for ray-finned fishes).

**Supplementary Table S2.** Regional indicator components by group. **Group-specific IndVal, specificity and fidelity for the taxonomic assignments displayed in Fig. 4C.** Each selected taxon–marker entry is evaluated in transition (nine transit samples), offshore (46 FPSO samples for 12S; 45 for 16S), and coast (eight samples from Urca and Ilhas Cagarras). An asterisk marks the associated group displayed in Fig. 4C; values for the other groups are descriptive comparisons, not additional significance tests. The p-value column reports the original permutation test of the maximum IndVal across the three groups (999 unrestricted sample-label permutations), with the maximizing group selected again in every permutation.

| Taxonomic assignment | Rank | Marker | Group | Detections / samples | Specificity A | Fidelity B | IndVal | Unadjusted p (max IndVal) |
| --- | --- | --- | --- | --- | --- | --- | --- | --- |
| Dactylopterus volitans | species | 16S | Transition * | 4/9 | 1 | 0.444444 | 44.44 | 0.002 |
| Dactylopterus volitans | species | 16S | Offshore | 0/45 | 0 | 0 | 0.00 | — |
| Dactylopterus volitans | species | 16S | Coast | 0/8 | 0 | 0 | 0.00 | — |
| Micropogonias | genus | 16S | Transition * | 6/9 | 0.660584 | 0.666667 | 44.04 | 0.006 |
| Micropogonias | genus | 16S | Offshore | 1/45 | 6.21782e-06 | 0.0222222 | 1.38e-05 | — |
| Micropogonias | genus | 16S | Coast | 1/8 | 0.33941 | 0.125 | 4.24 | — |
| Orthopristis | genus | 16S | Transition * | 3/9 | 1 | 0.333333 | 33.33 | 0.003 |
| Orthopristis | genus | 16S | Offshore | 0/45 | 0 | 0 | 0.00 | — |
| Orthopristis | genus | 16S | Coast | 0/8 | 0 | 0 | 0.00 | — |
| Harengula | genus | 16S | Transition * | 3/9 | 1 | 0.333333 | 33.33 | 0.005 |
| Harengula | genus | 16S | Offshore | 0/45 | 0 | 0 | 0.00 | — |
| Harengula | genus | 16S | Coast | 0/8 | 0 | 0 | 0.00 | — |
| Antigonia | genus | 12S | Transition * | 3/9 | 0.958754 | 0.333333 | 31.96 | 0.035 |
| Antigonia | genus | 12S | Offshore | 5/46 | 0.0412457 | 0.108696 | 0.45 | — |
| Antigonia | genus | 12S | Coast | 0/8 | 0 | 0 | 0.00 | — |
| Lepidophanes guentheri | species | 12S | Transition | 2/9 | 7.75876e-05 | 0.222222 | 0.00172 | — |
| Lepidophanes guentheri | species | 12S | Offshore * | 26/46 | 0.999922 | 0.565217 | 56.52 | 0.026 |
| Lepidophanes guentheri | species | 12S | Coast | 0/8 | 0 | 0 | 0.00 | — |
| Scombridae | family | 12S | Transition | 4/9 | 0.0979954 | 0.444444 | 4.36 | — |
| Scombridae | family | 12S | Offshore * | 28/46 | 0.902005 | 0.608696 | 54.90 | 0.035 |
| Scombridae | family | 12S | Coast | 0/8 | 0 | 0 | 0.00 | — |
| Scombridae | family | 16S | Transition | 1/9 | 0.0511374 | 0.111111 | 0.57 | — |
| Scombridae | family | 16S | Offshore * | 24/45 | 0.948863 | 0.533333 | 50.61 | 0.014 |
| Scombridae | family | 16S | Coast | 0/8 | 0 | 0 | 0.00 | — |
| Aluterus | genus | 12S | Transition | 3/9 | 0.00244305 | 0.333333 | 0.08 | — |
| Aluterus | genus | 12S | Offshore * | 22/46 | 0.997555 | 0.478261 | 47.71 | 0.044 |
| Aluterus | genus | 12S | Coast | 1/8 | 2.09474e-06 | 0.125 | 2.62e-05 | — |
| Engraulis | genus | 12S | Transition | 2/9 | 0.000234144 | 0.222222 | 0.01 | — |
| Engraulis | genus | 12S | Offshore | 13/46 | 0.0246922 | 0.282609 | 0.70 | — |
| Engraulis | genus | 12S | Coast * | 8/8 | 0.975074 | 1 | 97.51 | 0.001 |
| Sparidae | family | 12S | Transition | 1/9 | 0.000972823 | 0.111111 | 0.01 | — |
| Sparidae | family | 12S | Offshore | 0/46 | 0 | 0 | 0.00 | — |
| Sparidae | family | 12S | Coast * | 6/8 | 0.999027 | 0.75 | 74.93 | 0.001 |
| Mugil | genus | 16S | Transition | 2/9 | 0.0124214 | 0.222222 | 0.28 | — |
| Mugil | genus | 16S | Offshore | 0/45 | 0 | 0 | 0.00 | — |
| Mugil | genus | 16S | Coast * | 6/8 | 0.987579 | 0.75 | 74.07 | 0.001 |
| Mugil | genus | 12S | Transition | 3/9 | 0.232163 | 0.333333 | 7.74 | — |
| Mugil | genus | 12S | Offshore | 11/46 | 0.00275593 | 0.23913 | 0.07 | — |
| Mugil | genus | 12S | Coast * | 7/8 | 0.765081 | 0.875 | 66.94 | 0.002 |
| Brevoortia | genus | 12S | Transition | 6/9 | 0.279975 | 0.666667 | 18.66 | — |
| Brevoortia | genus | 12S | Offshore | 9/46 | 0.00454252 | 0.195652 | 0.09 | — |
| Brevoortia | genus | 12S | Coast * | 7/8 | 0.715483 | 0.875 | 62.60 | 0.006 |
| Megabalanus | genus | 16S | Transition | 0/9 | 0 | 0 | 0.00 | — |
| Megabalanus | genus | 16S | Offshore | 0/45 | 0 | 0 | 0.00 | — |
| Megabalanus | genus | 16S | Coast * | 5/8 | 1 | 0.625 | 62.50 | 0.001 |
| Engraulis anchoita | species | 16S | Transition | 0/9 | 0 | 0 | 0.00 | — |
| Engraulis anchoita | species | 16S | Offshore | 1/45 | 8.76589e-05 | 0.0222222 | 0.000195 | — |
| Engraulis anchoita | species | 16S | Coast * | 4/8 | 0.999912 | 0.5 | 50.00 | 0.001 |
| Paracentrotus | genus | 16S | Transition | 0/9 | 0 | 0 | 0.00 | — |
| Paracentrotus | genus | 16S | Offshore | 0/45 | 0 | 0 | 0.00 | — |
| Paracentrotus | genus | 16S | Coast * | 3/8 | 1 | 0.375 | 37.50 | 0.005 |

### Supplementary Data files — sampling, laboratory protocols, regional matrices and processing statistics

#### Curated supplementary matrices and final processing endpoints

The regional matrices contain all 64 regional biological samples and 11 Opentrons technical replicates, identified separately in the sample key; technical replicates are not independent biological samples.

Supplementary Data 4a–b contain author-curated derivatives of the taxonomic database count matrices. Supplementary Data 5a–b retain the database OTU identifiers, sequences and counts, with the five affected taxonomic annotations harmonized to the curated assignments. No OTUs were merged and no database records were changed. Supplementary Data 5a–b contain the OTU matrices used for downstream analyses: 212 OTUs for 12S and 241 for 16S, with 75 library columns each. Sequence-specific review excluded nine additional 12S sequences (107,893 reads) and eight additional 16S sequences (182,810 reads) as probable contaminants. The exclusion table records their taxon names, taxids and counts; no blanket taxonomic exclusion was applied.

Supplementary Data 3a–b report otus_final, reads_final and pct_reads_final for the OTU matrices, retain the supplied pre-additional-exclusion endpoints in separately named columns, and list excluded OTUs and reads per library. Negative controls PR347D, PR406D and PR534D have blank final fields; PR362 16S has 85 input pairs and zero final OTUs and reads.

The taxonomic and OTU matrices derive from different analysis runs and need not have identical totals. The regional 16S OTU snapshot contains 2,803 Mammalia reads absent from the taxonomic matrix; these reads were considered probable contamination.

Supplementary Data 1 provides sampling codes, coordinates and filtered water volumes; Supplementary Data 2 provides primer sequences, PCR reagents, thermocycling conditions, purification and normalization. The Supplementary Data archive contains six numbered TSV supplements (Data 4a–b, 5a–b and 3a–b), two method documents (Data 1 and Data 2), a data dictionary, a library key, an endpoint check, an exclusion log, a taxonomic-curation log and correction map, the updated Supplementary Table S1–S2 data, and a manifest.

- Data 1 — Sampling codes, coordinates and filtered water volumes (64 samples)
- Data 2 — PCR reagents and thermocycling conditions
- Data 3a — 12S processing statistics (78 libraries, including controls)
- Data 3b — 16S processing statistics (78 libraries, including controls)
- Data 4a — 12S taxonomic matrix (103 taxa × 75 libraries)
- Data 4b — 16S taxonomic matrix (131 taxa × 75 libraries)
- Data 5a — 12S OTU matrix (212 OTUs × 75 libraries)
- Data 5b — 16S OTU matrix (241 OTUs × 75 libraries)
- Data dictionary, source provenance and limitations
- Library-to-database key and biological/technical status
- Per-library verification of final processing endpoints and contaminant exclusions
- Author-identified probable contaminant sequences and exclusion counts
- Snapshot manifest and source hashes
- Supplementary Table S1 — class-composition data
- Supplementary Table S2 — indicator-component data
- Author taxonomic-curation log
- Taxonomic-curation correction map

