## Supplementary material for "Autonomous sampling and eDNA metabarcoding for offshore biodiversity monitoring: a pilot study around an FPSO in the Southwestern Atlantic": taxonomic-curation-log-20261019: taxonomic-curation-log-20261019.md

# 2026-10-19

## Síntese do dia

Foi realizada uma curadoria biogeográfica das tabelas finais de táxons do preprint para identificar atribuições em nível de espécie incompatíveis com a distribuição conhecida na costa brasileira. Cinco espécies foram classificadas como prováveis falsos positivos geográficos e reatribuídas ao gênero, nível taxonômico imediatamente superior com ocorrência documentada no Brasil. Quando a tabela já continha uma linha para o gênero correspondente, as contagens foram somadas por biblioteca para evitar linhas taxonomicamente redundantes. A correção foi restrita às tabelas suplementares do preprint; as tabelas canônicas de resultados e as matrizes ecológicas derivadas não foram alteradas.

## Atividades realizadas

### Curadoria biogeográfica das identificações taxonômicas finais

As tabelas `docs/preprint/supplementary-data-4a-taxa-12S.tsv` e `docs/preprint/supplementary-data-4b-taxa-16S.tsv` foram inspecionadas em conjunto com as anotações taxonômicas enriquecidas por GBIF e WoRMS presentes nas tabelas canônicas `identified_taxa.with-API-info.decontaminated.tsv`. A ocorrência das espécies no Brasil foi confrontada com registros e sínteses disponíveis no GBIF, WoRMS, OBIS, FishBase, ICMBio/SALVE e literatura taxonômica e biogeográfica. Também foi verificado se o nível imediatamente superior, principalmente o gênero, possuía ocorrência documentada no país.

Foram reconhecidas cinco atribuições em nível de espécie cuja distribuição aceita não inclui a costa brasileira:

- no 12S, *Brevoortia tyrannus* e *Cubiceps paradoxus*;
- no 16S, *Exocoetus monocirrhus*, *Luidia magnifica* e *Paracentrotus lividus*.

Os resultados dessa avaliação, incluindo distribuição mundial, situação da coluna `has_occurrence_in_brazil`, evidências para ocorrência do gênero no Brasil e fontes consultadas, foram preservados em `docs/preprint/probable-false-positives-12S.tsv` e `docs/preprint/probable-false-positives-16S.tsv`.

A inspeção mostrou que a coluna derivada do GBIF não era suficiente, isoladamente, para definir plausibilidade geográfica. *Brevoortia tyrannus* e *Exocoetus monocirrhus* estavam marcadas como presentes no Brasil, embora suas distribuições aceitas sejam, respectivamente, o Atlântico ocidental norte e o Indo-Pacífico/Pacífico oriental. No caso de *E. monocirrhus*, a consulta ao GBIF encontrou apenas um lote brasileiro sem coordenadas e com a ressalva `TAXON_MATCH_HIGHERRANK`. Em contraste, *L. magnifica*, *P. lividus* e *C. paradoxus* não apresentaram ocorrência brasileira no GBIF, em concordância com a literatura consultada. Em todos os cinco casos, o gênero possui representantes registrados no Brasil, justificando uma atribuição conservadora nesse nível.

### Reatribuição ao gênero e consolidação das leituras

Em `docs/preprint/supplementary-data-4a-taxa-12S.tsv`, a única leitura de *Brevoortia tyrannus* (taxon_id 110), observada em PR348D, foi somada à linha preexistente de *Brevoortia* (taxon_id 46; NCBI taxid 224706). A contagem desse gênero em PR348D passou de 650.397 para 650.398 leituras. Da mesma forma, a única leitura de *Cubiceps paradoxus* (taxon_id 341), observada em PR351D, foi incorporada à linha preexistente de *Cubiceps* (taxon_id 317; NCBI taxid 215400). As duas linhas em nível de espécie foram então removidas. A tabela passou de 105 para 103 táxons, mantendo inalterado o total de 106.441.553 leituras.

Em `docs/preprint/supplementary-data-4b-taxa-16S.tsv`, as 75 leituras de *Exocoetus monocirrhus* (taxon_id 265), restritas a PR364D, foram somadas à linha preexistente de *Exocoetus* (taxon_id 374; NCBI taxid 143331). A contagem do gênero nessa biblioteca passou de 662.434 para 662.509 leituras, e a linha da espécie foi removida. A identificação de *Exocoetus volitans* (taxon_id 407) foi mantida em nível de espécie por possuir ocorrência documentada no Brasil.

Como ainda não havia linhas de gênero correspondentes na tabela de 16S, *Paracentrotus lividus* (taxon_id 284) foi reescrita como *Paracentrotus*, com rank `genus`, NCBI taxid 7655 e linhagem truncada em `g__Paracentrotus`. As 6.994 leituras permaneceram distribuídas entre PR341D (2.221), PR343D (4.289) e PR345D (484). De modo equivalente, *Luidia magnifica* (taxon_id 409) foi reescrita como *Luidia*, com rank `genus`, NCBI taxid 7591 e linhagem truncada em `g__Luidia`. Suas 21.153 leituras foram preservadas nas oito bibliotecas originalmente positivas: PR359 (11), PR368 (29), PR368D (23), PR385D (2.540), PR388D (13.709), PR390D (140), PR393D (4.568) e PR396D (133).

Após as alterações, a tabela de 16S passou de 132 para 131 táxons, mantendo o total de 25.586.779 leituras. A validação final confirmou a ausência das cinco identificações em nível de espécie, a conservação das contagens por biblioteca, a manutenção dos totais globais e a inexistência de linhas duplicadas para os gêneros envolvidos.

## Pendências e próximos passos

- Manter explícito nos métodos do preprint que identificações geograficamente incompatíveis foram rebaixadas ao nível taxonômico superior com ocorrência comprovada no Brasil.
- Avaliar se tabelas, figuras ou textos derivados diretamente dos suplementos precisam ser regenerados após a redução de 105 para 103 táxons no 12S e de 132 para 131 táxons no 16S.
- Caso a mesma curadoria seja aplicada futuramente às tabelas canônicas e às matrizes ecológicas, repetir as análises derivadas e documentar separadamente seus impactos.
