## supplementary-data-guide for "Autonomous sampling and eDNA metabarcoding for offshore biodiversity monitoring: a pilot study around an FPSO in the Southwestern Atlantic": supplementary-data-guide.md

# ESP–offshore supplementary data: working package

**Status: author-curated taxonomic derivatives (2026-10-19); OTU counts and processing endpoints unchanged.**
On 2026-10-14 at 17:27, João Ferreira identified the 17 additional sequences in his attachments as probable contaminants and requested the same database matrices used for downstream ecological analyses. Data 4a–b were replaced on 2026-10-19 by the author's curated taxonomic tables (103 12S and 131 16S taxa). Data 5 keeps the same 212 and 241 OTU rows, sequences and counts; only their taxonomic annotation fields were harmonized to the curated names. Data 3 reports the retained OTU endpoints. No database records were changed. The original attachments remain intact as provenance, not the final analytical matrices.

## File inventory

| Supplement | File | Contents |
|---|---|---|
| Data 1 | supplementary-data-1-sampling.md | Sampling codes, coordinates and filtered volumes; 64 environmental samples |
| Data 2 | supplementary-data-2-pcr-thermocycling.md | Primers, reagents, thermocycling, purification and normalization |
| Data 2 Word | supplementary-data-2-pcr-thermocycling.docx | Latest author-approved methods and confirmed BSA quantities |
| Data 3a | supplementary-data-3a-processing-12S.tsv | Processing statistics with database-aligned final endpoints, 78 libraries including three negative controls |
| Data 3b | supplementary-data-3b-processing-16S.tsv | Processing statistics with database-aligned final endpoints, 78 libraries including three negative controls |
| Data 4a | supplementary-data-4a-taxa-12S.tsv | 103 curated taxonomic assignment taxa × 75 libraries |
| Data 4b | supplementary-data-4b-taxa-16S.tsv | 131 curated taxonomic assignment taxa × 75 libraries (PR362 explicit zero column) |
| Data 5a | supplementary-data-5a-otus-12S.tsv | 212 OTUs × 75 libraries; database OTU IDs and sequences; curated assignment annotations |
| Data 5b | supplementary-data-5b-otus-16S.tsv | 241 OTUs × 75 libraries; database OTU IDs and sequences; curated assignment annotations |

Data numbering is separate from the manuscript's Supplementary Tables and Figures; this avoids renumbering the manuscript figures and tables. Data 4a–b, 5a–b and 3a–b are UTF-8 tab-separated text with one header row. Data 1 and Data 2 are UTF-8 Markdown documents containing the method tables. Original attachments are byte-identical in ../artefatos/; current Data 3 files are explicitly derived, with original endpoints retained in separate columns. supplementary-data-manifest.json records hashes, sizes, exact runs and source provenance.

## Matrix format and library identity

Data 4 has one row per curated taxonomic taxon; Data 5 has one row per database OTU, its feature_id, sequence_sha256, representative_sequence and curated assignment metadata. Both have 75 library columns (PR*), with unnormalized integer counts from the relevant database run. No new blanket taxonomic filtering was applied. On 2026-10-19, five author decisions were applied: 12S taxon 110 (Brevoortia tyrannus) was summed into 46 (Brevoortia), 341 (Cubiceps paradoxus) into 317 (Cubiceps); 16S 265 (Exocoetus monocirrhus) into 374 (Exocoetus); 284 (Paracentrotus lividus) and 409 (Luidia magnifica) were relabelled to their genera, retaining the local taxon IDs. Per-library totals are unchanged (106,441,553 12S reads; 25,586,779 16S reads).

The 75 libraries comprise 64 biological samples and 11 technical replicates processed using Opentrons. Technical replicates are not independent biological samples. supplementary-data-sample-map.tsv connects each library label to its database sample_code and sample_id and flags the 47 biological libraries in the local subset, using the existing station metadata. It is a key, not a substitute for the complete spatial/environmental metadata and analysis scripts already in the project.

Zeros indicate no retained count in the exported matrix, not proof of biological absence. Sparse database count tables are expanded using the registered library roster. In particular, PR362 (ESP-OT:PR362) has an explicit all-zero 16S column: João's attachment reports 85 input pairs, 77 non-chimeric sequences, two OTUs immediately after clustering, and zero final OTUs/reads. This explicit zero is supported by the new source, not inferred from a missing database record. PR362D is a different library and is retained separately.

PR347D, PR406D and PR534D are negative controls, not biological samples. They occur in Data 3, but were removed from the final matrices and are not added to Data 4 or Data 5 as zero columns. Empty final fields for these controls mean not applicable/not present in the final matrix, never zero.

## Data 4: taxonomic fields

- taxon_id: TodaBio internal identifier, not an NCBI identifier.
- ncbi_taxid: NCBI Taxonomy identifier in the curated author table; original identifiers are retained in the correction map.
- scientific_name, taxon_rank, taxon_lineage: curated assignment identity, rank and lineage; these are derived metadata, not necessarily the unchanged database assignment.
- PR* columns: retained reads per library and assignment taxon from biodiv.sample_taxon_count, runs 8 (12S) and 9 (16S).

Bins have mixed taxonomic ranks. A family/genus-level unresolved assignment is not the aggregate of all species below that taxon. Bin counts are not species richness, and sequence counts are not organism abundance. Marker matrices must not be added to infer species richness.

## Data 5: OTU fields and provenance

Data 5a and 5b retain the 212 and 241 OTU rows of the archived database exports in ../artefatos/*-database-before-author-matrix.tsv. Every feature_id, representative sequence, sequence hash and library count is unchanged; only the five taxonomic annotations listed above were harmonized to the curated names in Data 4. OTUs were not merged. The 9 additional 12S sequences (107,893 reads) and 8 additional 16S sequences (182,810 reads) in the supplied attachments remain excluded from Data 5, in agreement with the database and the author's stated contaminant decision.

feature_id is the internal database identifier; sequence_sha256 identifies the exact representative sequence; assignment_status, taxon_id, ncbi_taxid, scientific_name, taxon_rank and taxon_lineage describe curated assignment derivatives. Original assignments remain in the archived source and correction map; molecular feature IDs are unchanged. Unassigned entries remain unassigned; taxon_id and ncbi_taxid are different identifier systems. The three negative controls are absent and PR362 16S is retained as an explicit zero column.

The file supplementary-data-contaminant-exclusions.tsv records all 17 review IDs, exact sequences, hashes, counts by library, taxon names and taxids provided by João, and attribution to his 17:27 message. These taxonomic identifications are author-provided, not independent sequence reassignments by this assistant. Exclusions apply to these exact sequences, not entire classes, orders or families. Original matrices remain in ../artefatos/biodiv_file_*.ontarget.decontaminated.tsv as source evidence, despite their filenames not representing the final additional exclusions.

## Data 3: processing-statistics dictionary

Definitions supplied by João Ferreira, 2026-10-14:

| Column | Meaning |
|---|---|
| sample | Library identifier, e.g. PR341D |
| negative_control | yes for PR347D, PR406D and PR534D (osmosis-water controls); no otherwise |
| marker | Library assay: 12s-mifish or 16s-marver3 |
| input | Read pairs entering DADA2; not independently established here as the total sequencer output before any upstream processing |
| filtered | Pairs passing DADA2 quality and length filtering |
| pct_filtered | 100 × filtered / input |
| denoised | Pairs for which both reads were corrected by the DADA2 error model |
| pct_denoised | 100 × denoised / input |
| merged | Pairs whose corrected reads were merged into one sequence |
| pct_merged | 100 × merged / input |
| non_chimeric | Merged sequences retained after chimera removal |
| pct_non_chimeric | 100 × non_chimeric / input |
| otus_99pct | Number of OTUs per library immediately after 99% clustering, before length/target filtering and contaminant removal; this is an OTU count, not a read count |
| otus_final | Number of OTUs with at least one read in the database-aligned Data 5 matrix, after the additional contaminant exclusion; blank for removed negative controls |
| reads_final | Sum of retained sequence counts in those final OTUs; blank for removed controls; zero for a retained library with no final counts |
| pct_reads_final | 100 × reads_final / input; blank when reads_final is blank |

input, filtered and denoised count pairs; merged/non_chimeric and reads_final count merged/retained sequences. One retained merged sequence derives from a pair; do not silently multiply counts by two or call every stage individual raw reads. All percentages use input, not the immediately preceding step. The supplied files do not separate losses due to length filtering, target filtering and decontamination after clustering; the additional exclusion is documented from exact sequence counts, without inventing earlier intermediate counts.

## Explicit additional-exclusion columns in Data 3

- supplied_before_additional_exclusion_otus_final, supplied_before_additional_exclusion_reads_final and supplied_before_additional_exclusion_pct_reads_final preserve the original endpoint fields exactly as supplied.
- additional_contaminant_otus_removed counts the excluded sequences with reads in that library; this is not the global number of unique excluded sequences.
- additional_contaminant_reads_removed sums the reads of those sequences in that library.
- otus_final, reads_final and pct_reads_final now describe the retained database OTU matrix; pct_reads_final = 100 × reads_final / input, rounded to two decimals.

All upstream processing fields are unchanged. For every retained library, the original OTU/read endpoint minus the explicit exclusions equals the current final endpoint. Controls retain blank final and exclusion fields; PR362 16S retains 85 input pairs and zero final OTUs/reads. The current processing files are derived, not byte-identical copies of the original attachments.

## Author clarification and scope

João identified the excluded sequences as Sus scrofa, Bos taurus, Hominidae, Cavia, Hemiptera and Homo sapiens (12S), and Hominidae, Canis lupus, Homo sapiens and Primates (16S), with taxids and counts recorded in supplementary-data-contaminant-exclusions.tsv. Probable contamination is the author's stated reason for their absence from the database, not an unresolved import failure.

The separately documented 2,803 regional 16S Mammalia reads present in the database OTU snapshot but absent from its taxonomic matrix remain a distinct taxon-versus-OTU difference; 2,674 occur in the local subset. João previously attributed that difference to probable contamination. No blanket exclusion of Mammalia is inferred, and no database matrix was modified to force taxonomic and OTU totals to agree. Data 4 and Data 5 preserve their respective runs, rather than representing identical taxonomic aggregation steps.

## Verification and historical records

The current supplementary-data-crosscheck.tsv compares the final processing endpoints with the database OTU matrix for all 150 marker/library combinations, with all endpoints matching. The original-minus-final differences are fully accounted for by the 17 author-identified sequences. Data 4a–b now carry the author's curated tables. Downstream ecological results that depend on species-level membership were recalculated in the isolated review branch; see review/science/REPORT.md.

Earlier author-matrices audits, author-endpoint-check TSVs, new-otu-cells TSVs, otu-database-map TSVs and the 17-OTU review ZIP are historical evidence of the pre-exclusion comparison, not current final supplements. They remain preserved and are explicitly listed as historical_only in the manifest; their earlier statements about unknown taxonomy or cause were superseded by João's 17:27 clarification.

Historical database-export scripts retain their original filenames and numbering in the project results directory. The historical 2026-10-16 delivery was renumbered without changing scientific TSV bytes. This 2026-10-19 revision supersedes Data 4 and the five affected annotation rows in Data 5. Current manifest hashes, author-input byte checks and sequence/count invariants verify the curated delivery; the old numbering verifier does not validate the new metadata.

No database correction is authorized: any future change requires prior consultation with João. Access for this work is biob_reader in READ ONLY transactions.

Delivery snapshot: project-sources/resultados/renumber-first-call-20261016/ (merged principal). Original exports and provenance remain in project-sources/resultados/. The Supplementary Data section offers downloadable HTTP assets and a ZIP archive under the same access control as the article. Local source paths are retained separately for provenance, not used as download links. Asset filenames include a content hash; download labels retain the original filenames. This is a delivery snapshot of the verified TSVs, not a new database export. Browser interaction by the requester has not been tested.

## Additional method supplements (2026-10-16)

Data 1 and Data 2 supersede Supplementary Table S1.2 and Supplementary Table S2, respectively, at João Ferreira’s request. Data 4a–b now carry the author’s corrected taxonomic tables (2026-10-19); Data 5a–b keep all OTU rows, sequences and counts with harmonized annotation metadata; Data 3a–b are unchanged. Data 1 and Data 2 are method documents, not paired marker matrices, so a/b suffixes do not apply. Viewing URLs remain stable; downloadable filenames use the Data identifiers.

## Numbering by first manuscript callout (2026-10-16)

Data 1: sampling; Data 2: primers and laboratory protocols; Data 3a–b: processing statistics; Data 4a–b: taxonomic matrices; Data 5a–b: OTU matrices. The a/b suffixes identify 12S/16S for paired datasets. Original source files are preserved at their historical paths; delivered filenames and all manuscript callouts use this numbering.

## Taxonomic-curation provenance (2026-10-19)

The archive includes taxonomic-curation-log-20261019.md (author-supplied rationale), taxonomic-curation-map-20261019.tsv (old/new IDs, names, ranks, NCBI identifiers and moved counts), and updated Supplementary Table S1/S2 data. Genus-level decisions were applied without inferring another species. No new BLAST, sequencing pipeline or database modification was performed. Preserved analyses were checked for numerical-input equivalence; taxonomic endpoints affected by aggregation or species membership were selectively recalculated. Original estimators, permutation definitions and cohorts were retained. Analysis code/results, source baseline and merge records are archived in the canonical project under resultados/taxonomic-curation-merge-20261019/.

## Supplementary Data 2 author update (2026-10-19)

Laboratory tables were updated from Danielle Amaral's revised document (Google Drive ID 1F_utd5FsBokNtXG4iEOJI9Tgzy5-Uenl). Primer sequences and numbered references remain unchanged. MD and editable DOCX contain identical tables. Source consistency notes flag unresolved BSA units, reaction-volume totals and the combined primer-volume entry; values were preserved rather than inferred. Data 1 and Data 3–5 retain their previous bytes. Earlier dated packages are historical provenance, not the current Data 2 version.


## Data 2 concentration and notation clarification (2026-10-19)

Danielle confirmed Illumina DNA/RNA UD Indexes, Tagmentation; precombined i5/i7 stock concentration 5x and final concentration 1x in both indexing tables. Remaining N/A table entries were replaced with a dash (-). Other values and source-consistency notes remain unchanged. Original imported sources and older delivery assets remain historical.


## Data 2 eLabJournal reconciliation (2026-10-19)

At Danielle Amaral’s instruction, eLabJournal per-reaction volumes supersede the earlier Drive transcription. BSA is 1.25 µL in A/C; water is 4.75 µL in A, 11.5 µL in D and 7.5 µL in E. B explicitly lists 1.25 µL of each MiFish-U primer. All reaction sums match declared totals. The 10% batch excess is not part of per-reaction volumes. BSA final-amount units remain an attributed unresolved source issue. Illumina indices 5x/1x and table dashes remain unchanged. Corrected Drive file retains its ID and link; original sources/assets remain historical.


## Completed operator-approved revisions (2026-10-19)

Current Data 2 includes only the protocols used for the sequenced libraries; the unsuccessful 16 July repeat-indexing recipe is retained solely as historical evidence in resultados/operator-approved-edits-20261019/. BSA supplier/catalog and stock, used index-set catalog numbers, manual reaction assembly, OT-2 Clean-up 2 and the NovaSeq X Series 1.5B 300-cycle kit are specified. The current Drive DNA entries were preserved. Primer sequences, numbered references, first-PCR 12S program and confirmed DreamTaq indexing program were preserved. BSA final-amount discrepancy remains explicitly flagged. The article now includes filter code, no pre-extraction wash and the third field blank. Only the authorized methods and corresponding automation-scope wording changed; analytical data, analyses, figures, Data 1/3–5 and João manuscript remain unchanged.


## BSA quantities confirmed by Danielle (2026-10-19)

The author confirmed final BSA amounts of 25 µg in each 25 µL 12S/16S amplification reaction and 50 µg in the 50 µL indexing reaction. Stock 20 mg/mL and volumes 1.25/2.5 µL yield final concentration 1 mg/mL. This supersedes all earlier BSA-pending notes; there is no remaining BSA quantity discrepancy in current deliverables. Historical source records and earlier snapshots remain preserved.

## Authorized integration into the principal (2026-10-19)

João Ferreira authorized the complete merge at 17:30 BRT. This delivery combines the reviewed taxonomic derivatives with the latest author-approved laboratory methods, confirmed BSA quantities, reference revisions, sample/taxon terminology, Figure 1 and regional ordination figures. No database write was performed. Historical exports are retained as provenance; current Data 4a–b reproduce the author-corrected tables byte for byte. Data 5 changes annotation metadata only, not OTU identifiers, sequences or counts.
