## supplementary-data-2-pcr-thermocycling for "Autonomous sampling and eDNA metabarcoding for offshore biodiversity monitoring: a pilot study around an FPSO in the Southwestern Atlantic": supplementary-data-2-pcr-thermocycling.docx

### Supplementary Data 2. PCR reagents and thermocycling conditions

#### Primer and overhang sequences

| Target or adapter | Primer / direction | Sequence (5′–3′) | Reference |
| --- | --- | --- | --- |
| 12S rRNA | MiFish-U FW | GTCGGTAAAACTCGTGCCAGC | [1] |
| 12S rRNA | MiFish-U RV | CATAGTGGGGTATCTAATCCCAGTTTG | [1] |
| 16S rRNA | MarVer3 FW | AGACGAGAAGACCCTRTG | [2] |
| 16S rRNA | MarVer3 RV | GGATTGCGCTGTTATCCC | [2] |
| Illumina overhang | FW | TCGTCGGCAGCGTCAGATGTGTATAAGAGACAG | [3] |
| Illumina overhang | RV | GTCTCGTGGGCTCGGAGATGTGTATAAGAGACAG | [3] |

**Sequence notation.** FW, forward; RV, reverse. All sequences are written 5′–3′. The four locus-specific primer sequences exclude the Illumina overhang. For overhang-bearing oligonucleotides, the corresponding FW or RV overhang is appended to the 5′ end of the locus-specific primer (5′–overhang–locus-specific sequence–3′). R in MarVer3 FW denotes A or G (IUPAC). Overhangs are distinct from the i5/i7 index sequences.

#### A. 12S MiFish-U: first PCR

| Component | Stock concentration | Volume (µL) | Final concentration or amount |
| --- | --- | --- | --- |
| Platinum SuperFi II PCR Master Mix (Thermo Fisher Scientific) | 2× | 12.5 | 1× |
| MiFish-U forward, without overhang | 10 µM | 1.25 | 0.5 µM |
| MiFish-U reverse, without overhang | 10 µM | 1.25 | 0.5 µM |
| BSA (Thermo Scientific; B14) | 20 mg/ml | 1.25 | 25 µg |
| DNA template | Approximately 5 ng/µL | 4 | Approximately 20 ng |
| Nuclease-free water | - | 4.75 | - |
| Total | - | 25 | - |

**Thermocycling**

| Stage | Temperature (°C) | Duration | Cycles |
| --- | --- | --- | --- |
| Initial denaturation | 95 | 3 min | 1 |
| Touchdown: denaturation | 94 | 30 s | 13 |
| Touchdown: annealing | 69.5 initially; decrease by 1.5 per cycle | 30 s | 13 |
| Touchdown: extension | 72 | 90 s | 13 |
| Amplification: denaturation | 94 | 30 s | 25 |
| Amplification: annealing | 60 | 30 s | 25 |
| Amplification: extension | 72 | 45 s | 25 |
| Final extension | 72 | 10 min | 1 |
| Hold | 4 | Indefinite | 1X |

#### B. 12S: second PCR for addition of Illumina overhangs

| Component | Stock concentration | Volume (µL) | Final concentration or amount |
| --- | --- | --- | --- |
| Platinum II Hot-Start PCR Master Mix (Thermo Fisher Scientific) | 2× | 12.5 | 1X |
| Platinum GC Enhancer | 5X | 5 | 1X |
| MiFish-U forward, with overhang | 10µM | 1.25 | 0.5µM |
| MiFish-U reverse, with overhang | 10µM | 1.25 | 0.5µM |
| First PCR product | Sample-dependent (0.089–3.96 ng/µL) | 1 | Variable (0.089–3.96 ng) |
| Nuclease-free water | - | 4 | - |
| Total | - | 25 | - |

**Thermocycling**

| Stage | Temperature (°C) | Duration | Cycles |
| --- | --- | --- | --- |
| Initial denaturation | 94 | 2 min | 1 |
| Denaturation | 94 | 15 s | 18 |
| Annealing | 57 | 30 s | 18 |
| Extension | 68 | 15 s | 18 |
| Final extension | 68 | 1 min | 1 |
| Hold | 4 | Indefinite | 1X |

#### C. 16S MarVer3: amplification with Illumina overhangs

| Component | Stock concentration | Volume (µL) | Final concentration or amount |
| --- | --- | --- | --- |
| Platinum SuperFi II PCR Master Mix (Thermo Fisher Scientific) | 2× | 12.5 | 1× |
| MarVer3 forward, with overhang | 10 µM | 2.5 | 1 µM |
| MarVer3 reverse, with overhang | 10 µM | 2.5 | 1 µM |
| BSA (Thermo Scientific; B14) | 20 mg/ml | 1.25 | 25 µg |
| DNA template | Approximately 5 ng/µL | 6.25 | Approximately 30 ng |
| Total | - | 25 | - |

**Thermocycling**

| Stage | Temperature (°C) | Duration | Cycles |
| --- | --- | --- | --- |
| Initial denaturation | 95 | 3 min | 1 |
| Touchdown: denaturation | 94 | 30 s | 11 |
| Touchdown: annealing | 55 initially; decrease by 0.5 per cycle | 30 s | 11 |
| Touchdown: extension | 72 | 90 s | 11 |
| Amplification: denaturation | 94 | 30 s | 27 |
| Amplification: annealing | 49 | 30 s | 27 |
| Amplification: extension | 72 | 45 s | 27 |
| Final extension | 72 | 10 min | 1 |
| Hold | 4 | Indefinite | 1X |

#### D. Indexing PCR: 12S and 16S

| Component | Stock concentration | Volume (µL) | Final concentration or amount |
| --- | --- | --- | --- |
| DreamTaq PCR Master Mix (Thermo Fisher Scientific) | 2× | 25 | 1× |
| BSA (Thermo Scientific; B14) | 20 mg/ml | 2.5 | 50 µg |
| Illumina DNA/RNA UD Indexes, Tagmentation; precombined i5/i7 | 5x | 10 | 1x |
| Purified amplicon with overhang | Approximately 5 ng/µL | 1 | Approximately 5 ng |
| Nuclease-free water | - | 11.5 | - |
| Total | - | 50 | - |

**Thermocycling**

| Stage | Temperature (°C) | Duration | Cycles |
| --- | --- | --- | --- |
| Initial incubation | 68 | 3 min | 1 |
| Initial denaturation | 98 | 3 min | 1 |
| Denaturation | 98 | 45 s | 12 |
| Annealing | 62 | 30 s | 12 |
| Extension | 68 | 2 min | 12 |
| Final extension | 68 | 1 min | 1 |
| Hold | 10 | Indefinite | 1X |

Index sets C (20091658) and D (20091660) were used for manually prepared 12S libraries and set A (20091654) for manually prepared 16S libraries. The additional technical preparations used set D. Indexing reactions were assembled manually; only clean-up was automated on the Opentrons OT-2 using the Clean-up 2 protocol.

#### E. Purification and normalization

| Step | Reagent or assay | Ratio or target |
| --- | --- | --- |
| After amplification | Illumina Purification Beads | 0.8:1 beads:DNA solution |
| After indexing | Illumina Purification Beads | 1.2:1 beads:DNA solution |
| Library normalization | Qubit High Sensitivity quantification | 2 nM |

**Notes.** Consecutive denaturation, annealing and extension rows with the same cycle count form one repeated cycle. A dash (-) indicates a value not applicable.

**Sequencing.** Illumina NovaSeq X Plus with the Illumina NovaSeq X Series 1.5B Reagent Kit (300 cycles), paired-end 2 × 150 bp.

**BSA.** The stock concentration was 20 mg/mL. Each 25 µL amplification reaction (12S and 16S) contained 1.25 µL of stock, corresponding to 25 µg BSA; each 50 µL indexing reaction contained 2.5 µL of stock, corresponding to 50 µg BSA. Final BSA concentration was 1 mg/mL in all three reactions. Volumes are per reaction; 10% excess applies only to preparation of batch mixes.
