## supplementary-data-1-sampling for "Autonomous sampling and eDNA metabarcoding for offshore biodiversity monitoring: a pilot study around an FPSO in the Southwestern Atlantic": supplementary-data-1-sampling.md

# Supplementary Data 1. Sampling codes, coordinates and filtered water volumes

| Sample | Filter code | Sampling group | Sampling code | Latitude (°) | Longitude (°) | Filtered volume (mL) |
| --- | --- | --- | --- | --- | --- | --- |
| PR341 | PR341F | Cagarras | — | -23.032601 | -43.213750 | 632 |
| PR342 | PR342F | Cagarras | — | -23.032601 | -43.213751 | 587 |
| PR343 | PR343F | Cagarras | — | -23.032601 | -43.213752 | 498.8 |
| PR344 | PR344F | Cagarras | — | -23.032601 | -43.213753 | 586 |
| PR345 | PR345F | Cagarras | — | -23.032601 | -43.213754 | 719 |
| PR346 | PR346F | Cagarras | — | -23.032601 | -43.213755 | 593.7 |
| PR348 | PR348F | Urca | — | -22.932265 | -43.153344 | 297 |
| PR349 | PR349F | Urca | — | -22.931101 | -43.153060 | 120 |
| PR350 | PR350F | FPSO | EQ#1 | -25.472764 | -43.938947 | 1000 |
| PR351 | PR351F | FPSO | BB#40 | -25.472764 | -43.938947 | 1000 |
| PR352 | PR352F | FPSO | BB#43 | -25.472764 | -43.938947 | 1000 |
| PR353 | PR353F | FPSO | EQ#5 | -25.472174 | -43.935426 | 1000 |
| PR354 | PR354F | FPSO | BB#39 | -25.472174 | -43.935426 | 1000 |
| PR355 | PR355F | FPSO | BB#30 | -25.472174 | -43.935426 | 1000 |
| PR356 | PR356F | FPSO | EQ#2 | -25.474640 | -43.936001 | 1000 |
| PR357 | PR357F | FPSO | BB#21 | -25.474640 | -43.936001 | 1000 |
| PR358 | PR358F | FPSO | BB#23 | -25.474640 | -43.936001 | 1000 |
| PR359 | PR359F | FPSO | EQ#4 | -25.471054 | -43.938959 | 1000 |
| PR360 | PR360F | FPSO | BB#24 | -25.471054 | -43.938959 | 1000 |
| PR361 | PR361F | FPSO | BB#29 | -25.471054 | -43.938959 | 1000 |
| PR362 | PR362F | FPSO | EQ#9 | -25.472315 | -43.942338 | 1000 |
| PR363 | PR363F | FPSO | BB#31 | -25.472315 | -43.942338 | 1000 |
| PR364 | PR364F | FPSO | BB#51 | -25.472315 | -43.942338 | 1000 |
| PR365 | PR365F | FPSO | EQ#3 | -25.473624 | -43.940974 | 1000 |
| PR366 | PR366F | FPSO | BB#25 | -25.473624 | -43.940974 | 1000 |
| PR367 | PR367F | FPSO | BB#34 | -25.473624 | -43.940974 | 1000 |
| PR368 | PR368F | FPSO | EQ#8 | -25.475522 | -43.943865 | 1000 |
| PR369 | PR369F | FPSO | BB#26 | -25.475522 | -43.943865 | 1000 |
| PR370 | PR370F | FPSO | BB#35 | -25.475522 | -43.943865 | 1000 |
| PR371 | PR371F | FPSO | EQ#7 | -25.479801 | -43.938862 | 1000 |
| PR372 | PR372F | FPSO | BB#28 | -25.479801 | -43.938862 | 1000 |
| PR373 | PR373F | FPSO | BB#47 | -25.479801 | -43.938862 | 1000 |
| PR374 | PR374F | FPSO | EQ#6 | -25.475283 | -43.933904 | 1000 |
| PR375 | PR375F | FPSO | BB#27 | -25.475283 | -43.933904 | 1000 |
| PR376 | PR376F | FPSO | BB#38 | -25.475283 | -43.933904 | 1000 |
| PR377 | PR377F | FPSO | BB#20 | -25.478740 | -43.935357 | 1000 |
| PR378 | PR378F | FPSO | BB#18 | -25.477507 | -43.936792 | 1000 |
| PR379 | PR379F | FPSO | BB#46 | -25.482083 | -43.932054 | 1000 |
| PR380 | PR380F | FPSO | EQ#11 | -25.484598 | -43.938825 | 1000 |
| PR381 | PR381F | FPSO | BB#37 | -25.484598 | -43.938825 | 1000 |
| PR382 | PR382F | FPSO | BB#44 | -25.484598 | -43.938825 | 1000 |
| PR383 | PR383F | FPSO | BB#45 | -25.489105 | -43.938785 | 1000 |
| PR384 | PR384F | FPSO | BB#36 | -25.481863 | -43.945945 | 1000 |
| PR385 | PR385F | FPSO | BB#42 | -25.469047 | -43.945769 | 1000 |
| PR386 | PR386F | FPSO | BB#52 | -25.469047 | -43.945769 | 1000 |
| PR387 | PR387F | FPSO | EQ#10 | -25.466544 | -43.939002 | 1000 |
| PR388 | PR388F | FPSO | BB#22 | -25.466544 | -43.939002 | 1000 |
| PR389 | PR389F | FPSO | BB#33 | -25.466544 | -43.939002 | 1000 |
| PR390 | PR390F | FPSO | BB#41 | -25.461903 | -43.939081 | 1000 |
| PR391 | PR391F | FPSO | BB#49 | -25.461903 | -43.939081 | 1000 |
| PR392 | PR392F | FPSO | BB#50 | -25.461903 | -43.939081 | 1000 |
| PR393 | PR393F | FPSO | BB#32 | -25.468707 | -43.932144 | 1000 |
| PR394 | PR394F | FPSO | BB#48 | -25.468707 | -43.932144 | 1000 |
| PR395 | PR395F | FPSO | BB#19 | -25.478050 | -43.942449 | 1000 |
| PR396 | PR396F | FPSO | BB#16 | -25.476497 | -43.941749 | 1000 |
| PR397 | PR397F | Transit | TR#1 | -25.369002 | -43.851335 | 1000 |
| PR398 | PR398F | Transit | TR#2 | -24.918005 | -43.640519 | 1000 |
| PR399 | PR399F | Transit | TR#3 | -24.445930 | -43.531387 | 1000 |
| PR400 | PR400F | Transit | TR#4 | -23.983801 | -43.397213 | 1000 |
| PR401 | PR401F | Transit | TR#5 | -23.566254 | -43.270469 | 1000 |
| PR402 | PR402F | Transit | TR#6 | -23.266330 | -43.220879 | 1000 |
| PR403 | PR403F | Transit | TR#7 | -23.169354 | -43.205454 | 1000 |
| PR404 | PR404F | Transit | TR#8 | -23.102494 | -43.192110 | 436 |
| PR405 | PR405F | Transit | TR#9 | -23.074000 | -43.182064 | 483 |

Coordinates are expressed in decimal degrees; negative values indicate south latitude and west longitude. Sample codes identify individual sampling events (EQ, BB or TR). A dash indicates that an EQ, BB or TR code does not apply.

The table includes 64 environmental samples: 47 from the FPSO area, nine from transit, six from Cagarras and two from Urca. Additional technical library preparations did not involve additional water filtration.
